# Coenzyme A biosynthesis in *Bacillus subtilis*: Discovery of a novel precursor metabolite for salvage and its uptake system

**DOI:** 10.1101/2024.04.17.589903

**Authors:** Robert Warneke, Christina Herzberg, Moritz Klein, Christoph Elfmann, Josi Dittmann, Kirstin Feussner, Ivo Feussner, Jörg Stülke

**Affiliations:** Department of General Microbiology, Institute for Microbiology & Genetics and Göttingen Center for Molecular Biosciences, University of Göttingen, 37077 Göttingen, Germany; Department of Plant Biochemistry, Albrecht-von-Haller Institute and Göttingen Center for Molecular Biosciences, University of Göttingen, 37077 Göttingen, Germany; Service unit for Metabolomics and Lipidomics, Göttingen Center for Molecular Biosciences, University of Göttingen, 37077 Göttingen, Germany

**Author notes:** **For correspondence:** Department of General Microbiology, Georg-August-University Göttingen, Grisebachstr. 8, 37077, Göttingen, Germany.

**Keywords:** *Bacillus subtilis*, coenzyme A, vitamin B5, metabolism, salvage, transport proteins

## Abstract

The Gram-positive model bacterium *Bacillus subtilis* is used for many biotechnological applications, including the large-scale production of vitamins. For vitamin B5, a precursor for coenzyme A synthesis, there is so far no established fermentation process available, partly due to the incomplete knowledge on the metabolic pathways that involve this vitamin. In this study, we have elucidated the complete pathways for the biosynthesis pantothenate and coenzyme A in *B. subtilis*. We have identified the enzymes involved in the pathway and have identified a salvage pathway for coenzyme A acquisition that acts on complex medium even in the absence of pantothenate synthesis. This pathway requires rewiring of sulfur metabolism resulting in the expression of a cysteine transporter. In the salvage pathway, the bacteria import cysteinopantetheine, a novel naturally occurring metabolite, using the cystine transport system TcyJKLMN. This work lays the foundation for the development of effective processes for vitamin B5 production.

## Introduction

The metabolism of humans and animals involves many compounds that are essential because they cannot be produced by the own metabolic processes. These compounds have therefore to be acquired from food. In addition to many essential amino acids and fatty acids, this is also true for vitamins. Many vitamins are used as precursors for important cofactors such as nicotinamide adenine dinucleotide (NAD), flavin adenine dinucleotide (FAD), thiamine pyrophosphate, tetrahydrofolic acid, pyridoxal phosphate, and coenzyme A. They are required in many metabolic pathways such as energy metabolism, central carbon metabolism, amino acid metabolism, or fatty acid biosynthesis. Microorganisms and plants are able to produce the vitamins, whereas animals and humans need to consume them with their food. Vitamins are used for many applications, including feed, food, cosmetics, chemicals and pharmaceutics, and thus need to be produced on a large scale (1).

Vitamins can be produced by chemical synthesis or by biotechnological processes. Biotechnology is regarded as more sustainable, and a variety of microorganisms such as *Escherichia coli*, *Bacillus subtilis*, *Corynebacterium glutamicum*, *Ashbya gossypii* or baker’s yeast *Saccharomyces cerevisiae* are in use for such processes or targets for process development (2). An example for the replacement of a chemical by a biotechnological process is the production of vitamin B2 (riboflavin), a precursor for the flavin coenzymes FAD and FMN. In this case, the biotechnological process has replaced the chemical synthesis in as little as 15 years due to economic and sustainability benefits (3). We are interested in the physiology of the Gram-positive model bacterium *B. subtilis*. In addition to being one of the best-studied bacteria, *B. subtilis* is also a major workhorse in biotechnology for the production of proteins and vitamins and many other applications (3, 4, 5). Moreover, *B. subtilis* is the target of attempts to minimize the genome in order to get a comprehensive understanding for the component requirements of a living cell (6, 7, 8) and to use genome-minimized strains as platforms in biotechnological applications (9, 10, 11).

Coenzyme A is an essential cofactor that is involved in many metabolic pathways such as central carbon metabolism, amino acid metabolism, and fatty acid biosynthesis. This cofactor is synthesized from pantothenic acid which is also known as vitamin B5. *B. subtilis* is able to synthesize coenzyme A from two molecules of pyruvate, aspartate, and cysteine (see Fig. 1) with pantothenate being an intermediate of the pathway. The initial reactions from pyruvate to α-ketoisovalerate are shared with branched chain amino acid biosynthesis and are catalyzed by enzymes of this pathway. For the specific steps of pantothenate and coenzyme A biosynthesis, enzymes have been identified biochemically or based on the similarity to enzymes from other organisms for each reaction of the pathway (see Fig. 1). Even though both vitamin B5 and coenzyme A are very important metabolites, their metabolism is not fully understood in the biotechnological vitamin production platform *B. subtillis*. For the conversion of α-ketopantoate to pantoate and the phosphorylation of pantothenate, two enzymes each (YlbQ/YkpB and CoaA/CoaX, respectively) have been suggested, but their specific roles have not been clarified. Moreover, for the *panC* mutant lacking pantothenate synthase, it has been shown that such a mutant is viable on complex, but not on minimal medium, suggesting that intermediate(s) of coenzyme A biosynthesis can be taken up from complex medium. In addition, the *panB* and *panC* mutants are impaired in growth even on complex medium, and the *panC* mutant was reported to readily acquire suppressor mutations (12). This suggests the existence of other enzymes and/or means of synthesizing coenzyme A in *B. subtilis*.

**Figure 1:**
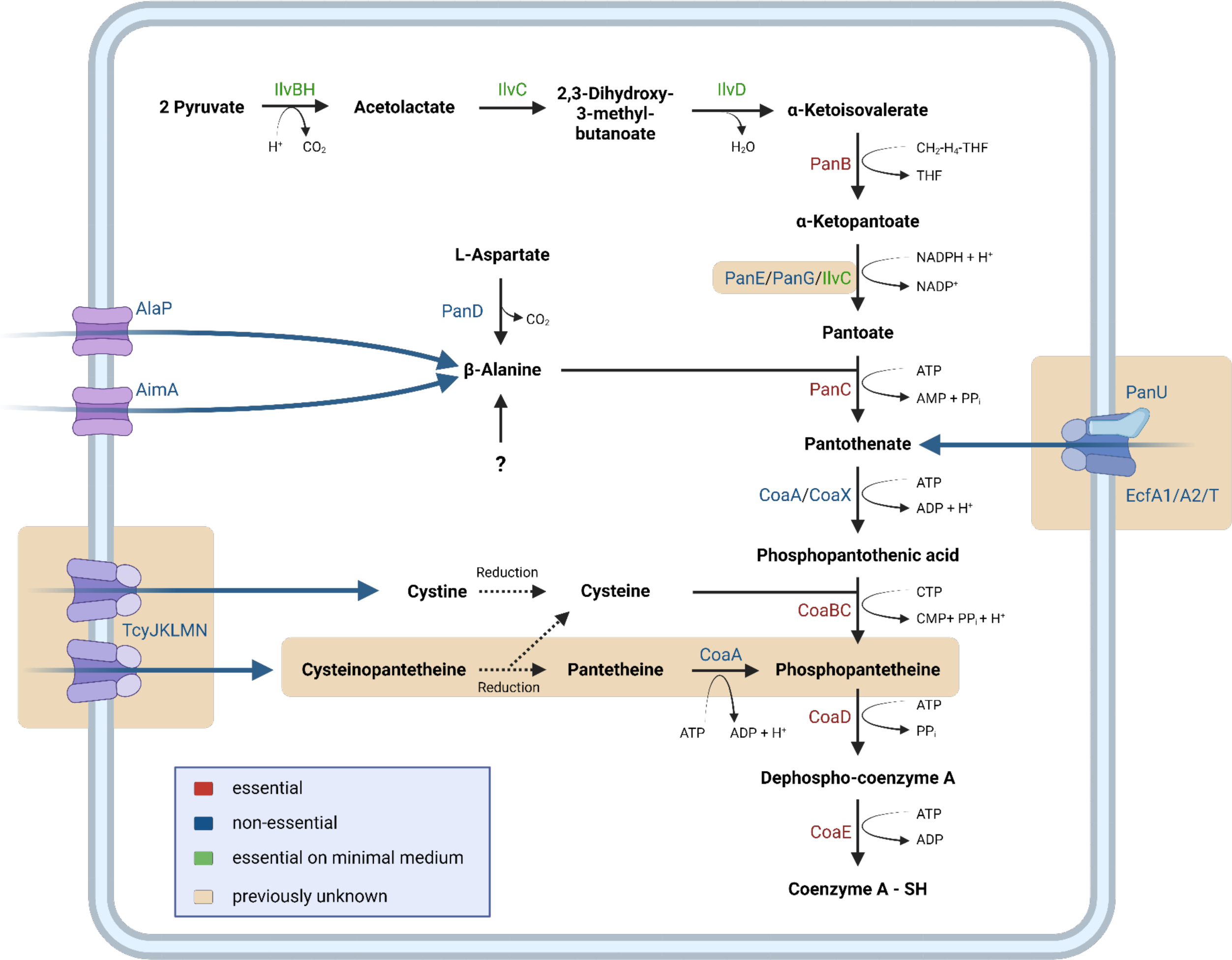
Schematic overview of the pathways for coenzyme A acquisition in *B. subtilis*. The biosynthetic and salvage pathways as well as the uptake systems for important precursors are shown. Enzymes and reactions identified in this study are highlighted in light brown.

*B. subtilis* is one of the organisms that are central for the development of the fermentative production of vitamin B5, to allow the switch from chemical to biotechnological synthesis. Using strains overexpressing enzymes of the pathway, up to 80 g/l of vitamin B5 could be produced (2). However, to optimize the biotechnological production of a metabolite, it is essential to comprehensively understand the complete biosynthetic pathway as well as alternative routes that feed into it. As discussed above, our knowledge of the B5 pathway in *B. subtilis* is still limited. We have recently studied two pathways that feed into B5 synthesis. For the *ilvBHC-leuABCD* operon that encodes enzymes for α-ketoisovalerate biosynthesis, we demonstrated that its expression is activated in the presence of preferred carbon sources by the pleiotropic transcription factor CcpA (13). Moreover, this operon is repressed in the presence of amino acids and activated under conditions of nitrogen limitation by the transcription factors CodY and TnrA, respectively (14, 15). In addition, we have recently studied the homeostasis of the precursor molecule β-alanine (16). This molecule can be taken up from the medium by the amino acid importers AimA and AlaP. Interestingly, intracellular accumulation of β-alanine is toxic for *B. subtilis*, and the molecule can be exported by the amino acid exporter AexA. The corresponding *aexA* gene is silent under most conditions and can only be expressed upon the acquisition of mutations that trigger DNA-binding activity of the transcription activator AerA (16).

In this work, we have studied the pathway leading to the formation of vitamin B5 and coenzyme A in *B. subtilis* in detail. We have identified three enzymes that contribute to the reduction of α-ketopantoate as well as an uptake system for vitamin B5, pantothenic acid. In addition to pantothenate, *B. subtilis* can also use pantetheine as a precursor for coenzyme A biosynthesis. Our work demonstrates that the two pantothenate kinases have shared and individual activities. The analysis of suppressor mutants that allow the growth of *panB* or *panC* mutants identified cysteinopantetheine, a novel metabolite, as an additional precursor that can be taken up by *B. subtilis* and feed into coenzyme A biosynthesis, bypassing the synthesis of vitamin B5. The use of cysteinopantetheine requires the expression of the CymR-repressed *snaA-tcyJKLMN-cmoOIJ-ribR-sndA-ytnM* operon to allow its uptake by the TcyJKLMN ABC transporter.

## Results

### Essential and conditionally essential enzymes in the biosynthesis of CoA

The pathway for CoA biosynthesis has not been completely elucidated in *B. subtilis* (see Fig. 1). Based on a large-scale attempt to construct mutants for all genes, the genes for the last three steps of CoA biosynthesis, i. e. *coaBC*, *coaD*, and *coaE*, are essential in *B. subtilis*, indicating that they are required for growth even on complex medium (12). However, the genes for the initial steps of the pathway to pantothenate could be deleted, but their fitness was impaired on complex medium and in some cases the mutants were unable to grow on minimal medium (see Fig. 2A). To test the role of these genes, we assayed growth of all mutants starting with the first dedicated enzyme of the pathway, α-ketoisovalerate hydroxymethyltransferase (PanB, see Fig. 1, Box 1) to the last non-essential enzyme, pantothenate kinase (CoaA, CoaX). As shown in Fig. 2B, the *panB* and *panC* mutants GP4401 and GP4402, respectively, were not viable on the complex medium used here, whereas growth of all other mutants was indistinguishable from that of the wild type strain 168. This indicates that the *panB* and *panC* genes are also essential for the growth of *B. subtilis*. For the *panB* mutant, the appearance of individual colonies is visible at the highest cell concentration. This suggests the formation of suppressor mutants (see below). Indeed, it has been reported that several mutants of the pathway tend to acquire suppressor mutations (12). In contrast, all mutants were viable on the medium supplemented with pantothenate. Thus, the *panB* and *panC* genes can be regarded as conditionally essential, as they are dispensable if the cells are provided with the product of the pathway. β-Alanine, a precursor for pantothenate synthesis, can be taken up from the environment or is synthesized by the aspartate decarboxylase PanD (16, 17). The *panD* mutant was viable both on complex and minimal medium (Fig. 2B), indicating that additional enzymes contribute to β-alanine synthesis. The *coaA* and *coaX* mutants grew indistinguishably from the wild type strain, as reported previously (18). However, we were unable to construct a *coaA coaX* double mutant, indicating that there is no additional pantothenate kinase in *B. subtilis*. Taken together, our findings indicate that pantothenate is an essential precursor for CoA biosynthesis, and that *B. subtilis* can take up this important intermediate from the environment.

**Figure 2:**
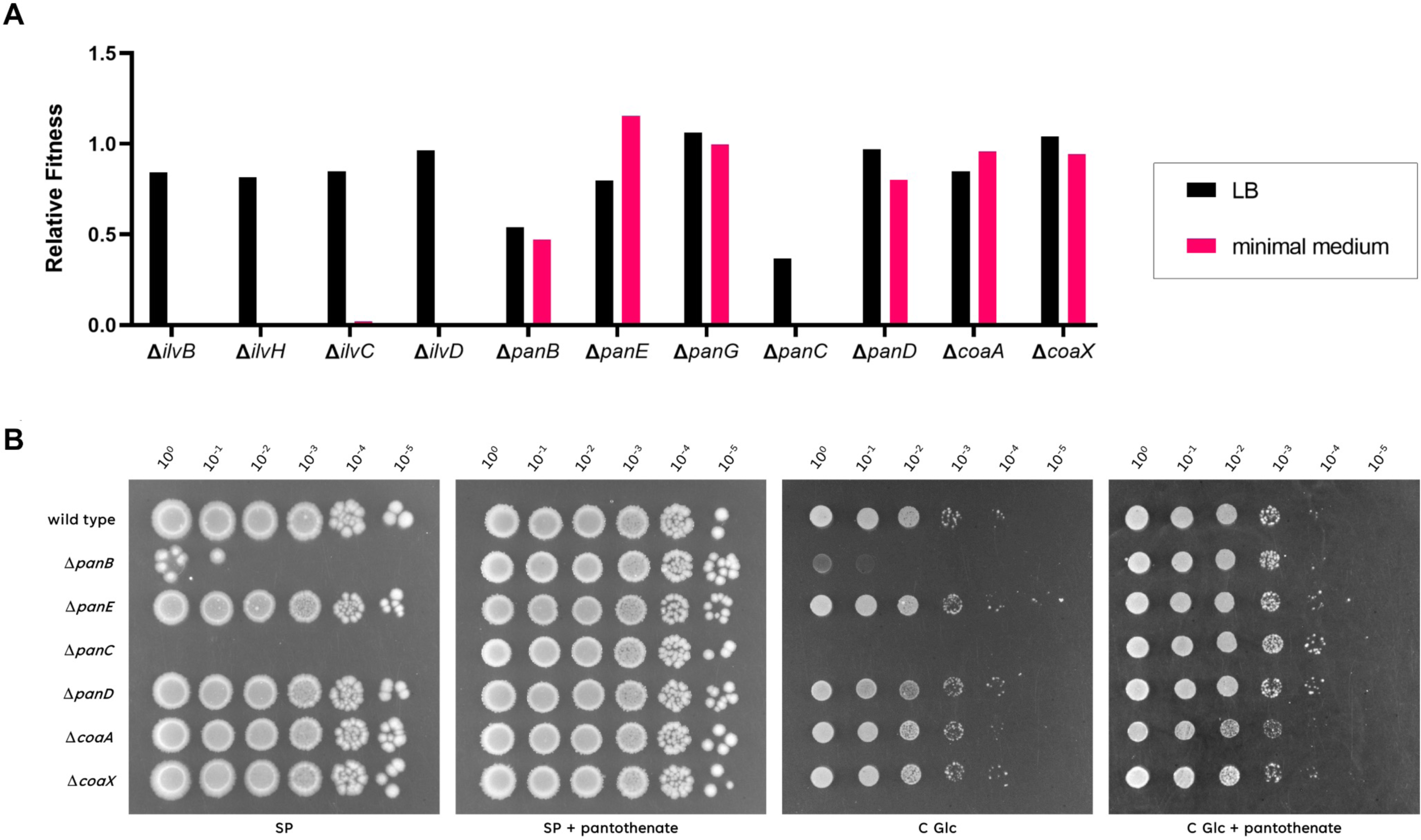
Essential and conditionally essential enzymes in the biosynthesis of coenzyme A. A, The relative fitness of mutant strains (12). B, The growth of the *B. subtilis* wild type was compared with the mutants Δ*panB* (GP4401), Δ*panE* (GP3384), Δ*panC* (GP4402), Δ*panD* (GP3382), Δ*coaA* (GP3380) and Δ*coaX* (GP3381). The cells were grown in sporulation medium (SP) supplemented with 1 mM pantothenate to an OD_600_ of 1.0, and serial dilutions (10-fold) were prepared. These samples were plated on SP or C-Glc minimal plates with or without added 1 mM pantothenate and incubated at 37°C for 48 h.

### Identification of enzymes that contribute to pantoate biosynthesis in *B. subtilis*

The vitamin B5 precursor pantoate is synthesized from α-ketoisovalerate, an intermediate in valine biosynthesis. In the first step, α-ketoisovalerate is hydroxymethylated to 2-ketopantoate. In the next step, 2-ketopantoate is reduced to pantoic acid by ketopantoate reductase PanE (2). *B. subtilis* encodes two members of the ketopantoate reductase family, YlbQ and YkpB. These proteins have so far not been characterized. In the UniProt database, the YlbQ protein is annotated as PanE; accordingly, the corresponding mutant GP3384 was included in our initial growth assay (see Fig. 2B). However, growth of the *panE* mutant was not impaired indicating that the second putative ketopantoate reductase YkpB (re-designated PanG, see Box 1) also contributes to ketopantoate reduction. To get insights into the roles of PanE and PanG in pantoate biosynthesis, we tested growth of single and double mutants on minimal medium (see Extended Data Fig. 1). All strains grew well indicating the presence of yet other enzymes that may contribute to pantoate formation. In *E. coli*, the ketol-acid reductoisomerase IlvC, which is primarily involved in branched-chain acid amino acid biosynthesis, is rather promiscuous and can act on a variety 2-ketoacids, including 2-ketopantoate (19). We thus considered the possibility that IlvC might be a third pantoate-forming enzyme in *B. subtilis*. Indeed, growth of the *panE panG ilvC* triple mutant GP4403 was severely impaired on complex medium in the absence of pantothenate, indicating that ketopantoate reduction is an essential function and that IlvC is the third ketopantoate reductase in *B. subtilis*.

### Identification of an uptake system for pantothenate

The results presented above demonstrate that the *panB* and *panC* mutants are auxotrophic for pantothenate (see Fig. 2B). Vitamins and cofactors are often transported by multimeric ECF-type transporters that consist of a transmembrane transport protein (YbaF in *B. subtilis*), two ATP-binding subunits (YbxA and YbaE), and substrate-specific binding proteins (20, 21). To test whether pantothenate is also taken up by an ECF transporter in *B. subtilis*, we compared the growth of the wild type strain 168, *panC* mutant GP4361 and the isogenic *panC ybaF* (*ecfT*) double mutant GP4700 (see Fig. 3A). While the wild type strain was able to grow in the absence of pantothenate, both strains lacking *panC* required the addition of pantothenate to the medium. If pantothenate uptake would require an ECF transporter, the double mutant GP4700 would be unable to grow, as was indeed observed at a low pantothenate concentration (50 µM). In contrast, the *panC ybaF* (*ecfT*) double mutant was viable at a high pantothenate concentration (1 mM). As a control, we also tested the growth of the *ybaF* (*ecfT*) single mutant GP4703, which was viable in the absence of pantothenate. This indicates that the phenotype at the low pantothenate concentration resulted from the combined absence of pantothenate biosynthesis and transport. Taken together, these data indicate that the ECF transport system is responsible for high affinity pantothenate transport, and that an additional uptake system can transport the intermediate with low affinity.

**Figure 3:**
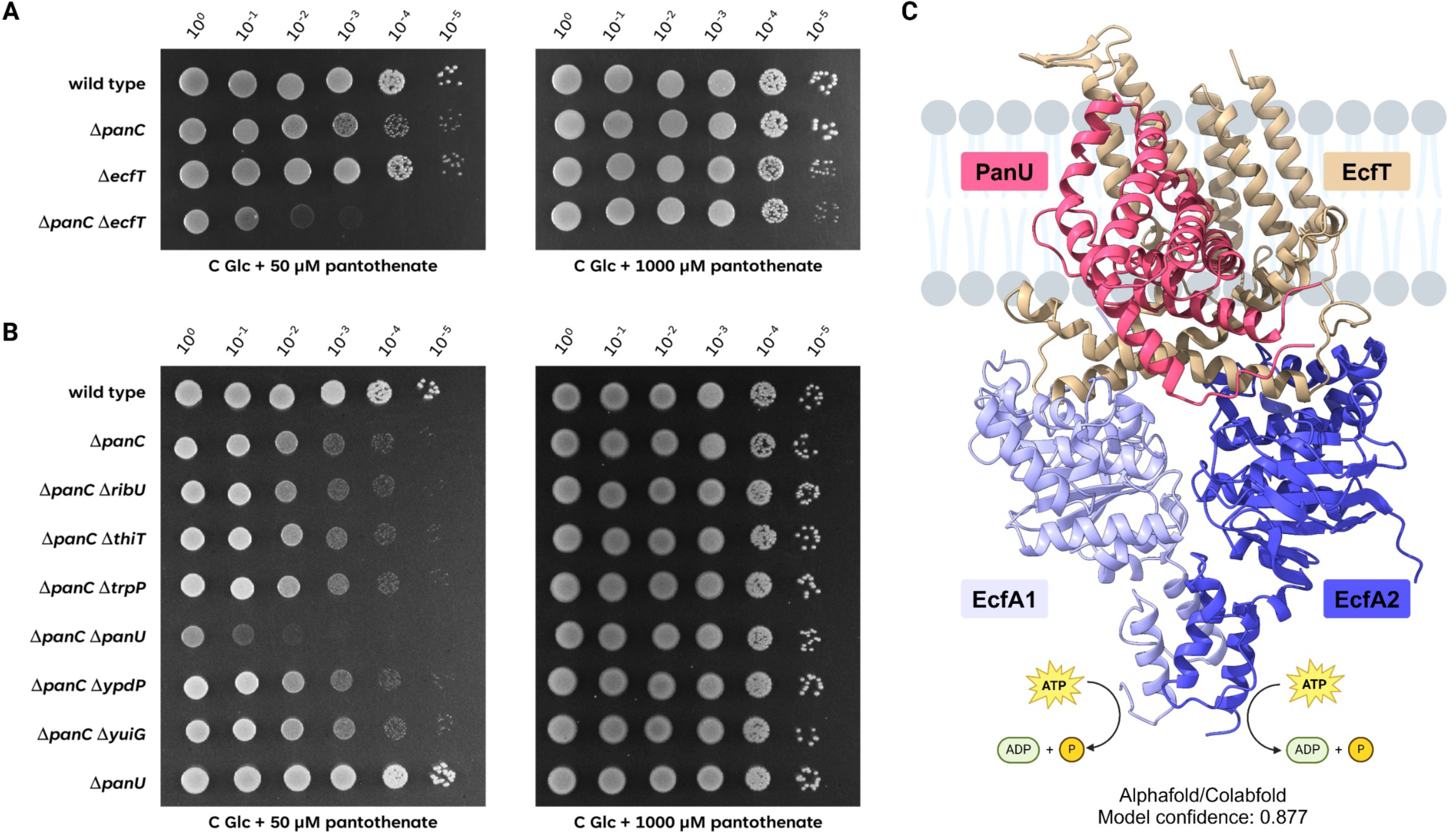
Pantothenate is imported via the ECF transporter EcfA1/A2/T with the S component PanU. A, The growth of the *B. subtilis* wild type was compared with the mutants Δ*panC* (GP4361), Δ*ecfT* (GP4703), and the double mutant Δ*panC* Δ*ecfT* (GP4700). The cells were grown in C glucose minimal medium supplemented with 1 mM pantothenate to an OD_600_ of 1.0, and serial dilutions (10-fold) were prepared. These samples were plated on C Glc minimal medium supplemented with 50 µM or 1 mM pantothenate and incubated at 37°C for 48 h. B, The growth of the Δ*panC* mutant (GP4361) was compared with the isogenic Δ*ribU* (GP4694), Δ*thiT* (GP4695), Δ*trpP* (GP4696), Δ*panU* (GP4697), Δ*ypdP* (GP4698) and Δ*yuiG* (GP4699) mutants, as well as the Δ*panU* (GP4704) strain. The cells were grown in C glucose minimal medium supplemented with 1 mM pantothenate to an OD_600_ of 1.0, and serial dilutions (10-fold) were prepared. These samples were plated on C Glc minimal medium supplemented with 50 µM or 1 mM pantothenate and incubated at 37°C for 48 h. C, ColabFold/AlphaFold predicted model of the complex of EcfT (beige) – EcfA1 (light blue) – EcfA2 (dark blue) – PanU (magenta). The model was placed on a schematic membrane (gray) based on position of the transmembrane helices of EcfT and PanU. As indicated below the structure, the model confidence (iPTM score) of the calculated model is 0.877.

As mentioned above, the ECF transporters consist of three general components and a substrate-specific binding protein, the S protein. To identify the S protein required for high-affinity pantothenate transport, we deleted all genes encoding the S proteins of *B. subtilis* in the background of the *panC* mutant and tested their growth in the absence and presence of pantothenate (Fig. 3B). With the exception of the *panC yhfU* (*panU*) double mutant GP4697, all strains were viable if pantothenate was present in the medium. The deletion of the *yhfU* (*panU*) gene by itself had no impact on the viability of *B. subtilis*, since the single *yhfU* (*panU*) mutant GP4704 grew well in the absence of pantothenate, reflecting its ability to synthesize the intermediate.

Taken together, our results demonstrate that pantothenate can be imported by *B. subtilis* with high affinity using an ECF transporter with YhfU as the substrate-specific S protein. Therefore, we rename the general ECF transporter proteins EcfA1 and EcfA2 (the ATPase components, previously YbxA and YbaE, respectively, see Box 1) and EcfT (the membrane-spanning transporter component, previously YbaT), and the pantothenate-binding S protein PanU (for <u>pan</u>tothenate <u>u</u>ptake, previously YhfU).

We used ColabFold to predict a structural atomic model of the EcfA1/ EcfA2/EcfT/ PanU complex. The highest ranking model had a high local accuracy as indicated by the predicted local distance difference test (pLDDT) value of 85.8. It also featured high pTM (0.867) and ipTM (0.877) scores which indicate high confidence in the folding of the protein subunits and the complex interfaces, respectively. The model (see Fig. 3C) predicts that EcfT and PanU form a transmembrane complex for pantothenate binding and transport, whereas the ATP-binding EcfA1 and EcfA2 subunits are bound to this complex via cytoplasmic coupling helices of EcfT. The EcfA1 and EcfA2 subunits additionally interact with each other via their C-terminal domains (see http://www.subtiwiki.uni-goettingen.de/v4/predictedComplex?id=173 for an interactive presentation of the complex). The model is in excellent agreement with the cryo-electron microscopic structure of the folate ECT transporter of *Lactobacillus delbrückii* (22).

### A CoaA- and pantetheine-dependent salvage pathway

In *E. coli*, a pantothenate-independent pathway for coenzyme A biosynthesis was discovered (23). In this bypass, pantethine and/or pantetheine are taken up, and pantetheine is directly phosphorylated to phosphopantetheine. This reaction is catalyzed by the single pantothenate kinase of *E. coli*, CoaA (23). Interestingly, many bacteria possess either CoaA or CoaX, and only few bacteria including *B. subtilis* have both enzymes. *E. coli* encodes only the type I pantothenate kinase CoaA, and this enzyme has a second activity as a pantetheine kinase. To test whether this bypass is also active in *B. subtilis*, we assayed the growth of the *panC* mutant GP4379 and the isogenic double mutants *panC coaA* (GP4652) and *panC coaX* (GP4653), in the presence of pantethine (see Extended Data Fig. 2A). In *E. coli*, pantethine is taken up and reduced to pantetheine (23). The *panC* mutant was viable indicating that *B. subtilis* is also able to use pantetheine as a precursor for coenzyme A biosynthesis. Of the two kinases, only CoaA was required for the utilization of pantetheine, whereas the loss of CoaX had no effect. This indicates that CoaA has pantetheine kinase activity in addition to its activity as a pantothenate kinase, whereas CoaX only has the latter activity. This is in good agreement with the activity of *E. coli* CoaA, which also has both activities. The CoaA enzymes from *E. coli* and *B. subtilis* share 50% identical residues, which may explain why the proteins have the same biochemical activities.

The existence of this salvage pathway is further supported by the investigation of a *coaBC* mutant. The *coaBC* (previously *yloI*) gene of *B. subtilis* is essential for growth (12, 24). Based on the results described above, pantetheine should rescue the *coaBC* mutant. Indeed, we were able to delete the *coaBC* gene, provided that pantetheine was present in the growth medium (see Extended Data Fig. 2B). The resulting *coaBC* mutant GP4670 was only viable in the presence of pantetheine. This observation supports the proposed pantetheine and CoaA-dependent salvage route to coenzyme A in *B. subtilis* (see Fig. 1).

### Isolation of suppressor mutants that can grow in the absence of internally synthesized pantothenate

As mentioned above, we observed the appearance of suppressor mutants when we cultivated the *panB* mutant in the absence of pantothenate. Similarly, suppressor formation was also observed for the *panC* mutant. Two suppressor mutants were obtained from the *panB* mutant, and one from the *panC* mutant. The genomes of all three suppressor mutants were analyzed by whole genome sequencing (see Fig. 4A). The two strains derived from the *panB* mutant both had an identical point mutation in the *cysK* gene, encoding the cysteine synthase, the last enzyme of cysteine biosynthesis (25). The mutation resulted in the formation of a truncated CysK protein (G223 Stopp). Interestingly, strain GP4450 derived from the *panC* mutant also carried a mutation that may affect cysteine metabolism. In this case, the *cymR* gene, encoding the transcriptional repressor of a regulon involved in cysteine metabolism (26), was affected, resulting in a CymR protein with an amino acid substitution of Leu-49 to Thr. This substitution affects the DNA-binding helix-turn-helix motif of CymR (27), thus resulting in loss of CymR-mediated repression. Two of the analyzed suppressor mutants had additional mutations (see Fig. 4A).

**Figure 4:**
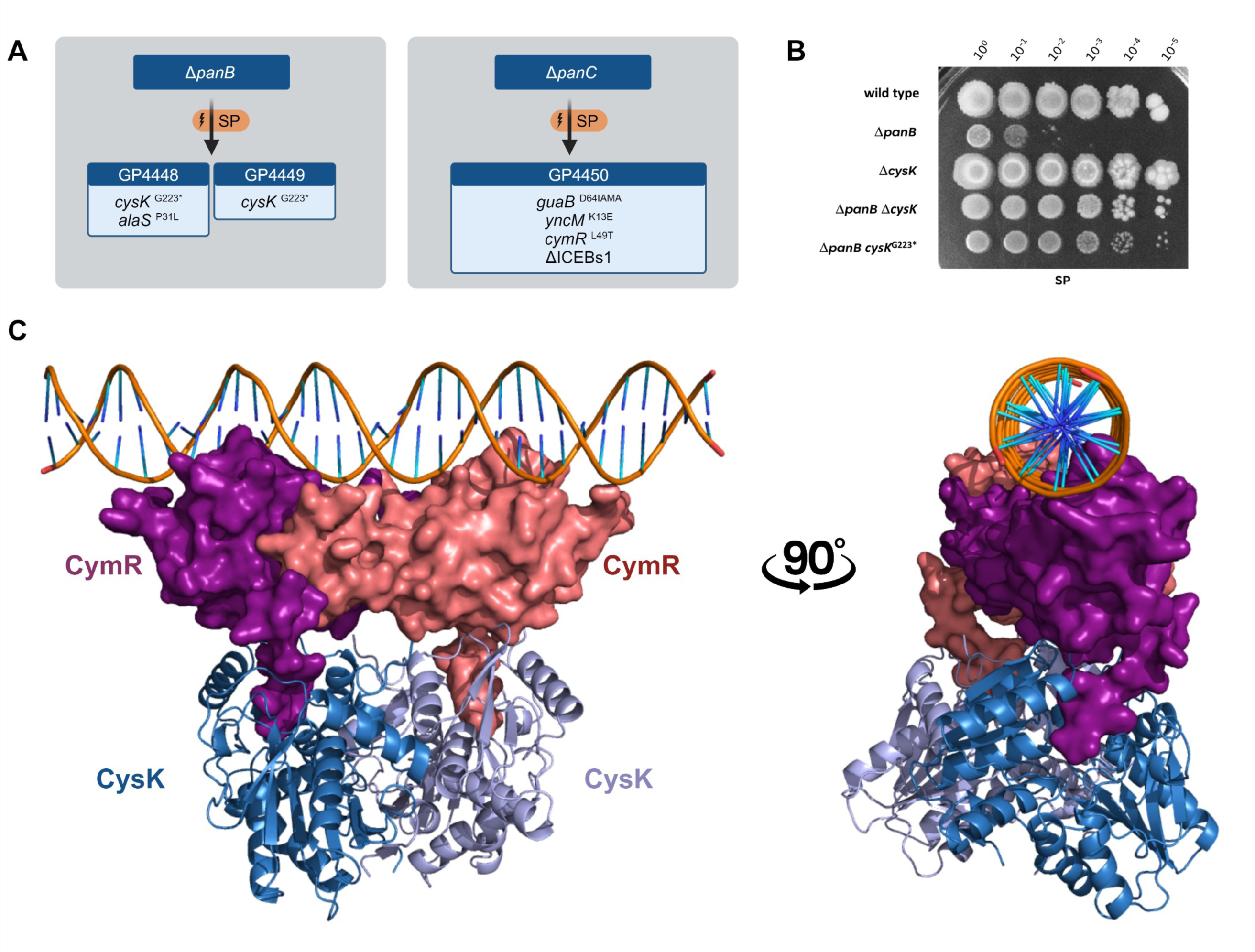
Suppressors of mutants deficient in pantothenate biosynthesis inactivate the CymR-CysK repressor complex. A, Evolutionary trajectory of the Δ*panB and* Δ*panC* mutants on sporulation medium (SP) without supplemented pantothenate. B, The growth of the *B. subtilis* wild type was compared with the mutants Δ*panB* (GP3383), Δ*cysK* (GP4404), Δ*panB* Δ*cysK* (GP3343) and the suppressor mutant Δ*panB cysK*^G223*^ (GP3397). The cells were grown in sporulation medium (SP) supplemented with 1 mM pantothenate to an OD_600_ of 1.0, and serial dilutions (10-fold) were prepared. These samples were plated on SP plates and incubated at 37°C for 48 h. C, ColabFold/AlphaFold predicted model of the complex of the CymR dimer (red, purple) with the CysK dimer (lightblue, darkblue) bound to the 5’UTR of the *cysK* gene (modelled with HDOCK).

To test whether the *cysK* or *cymR* mutations are sufficient to allow growth of the *panB*/ *panC* mutants in the absence of added pantothenate, we deleted the *cysK* and *cymR* genes in the *panB* and *panC* mutants, respectively, and tested the growth of the double mutants. As shown in Fig. 4B, the *panB cysK* double mutant GP4124 was able to grow without added pantothenate. Similarly, the deletion of *cymR* allowed growth of the *panC* mutant (see Fig. 5A). These results demonstrate that the inactivation of either *cysK* or *cymR* is necessary and sufficient to suppress the pantothenate deficiency of the *panB* and *panC* mutants.

**Figure 5:**
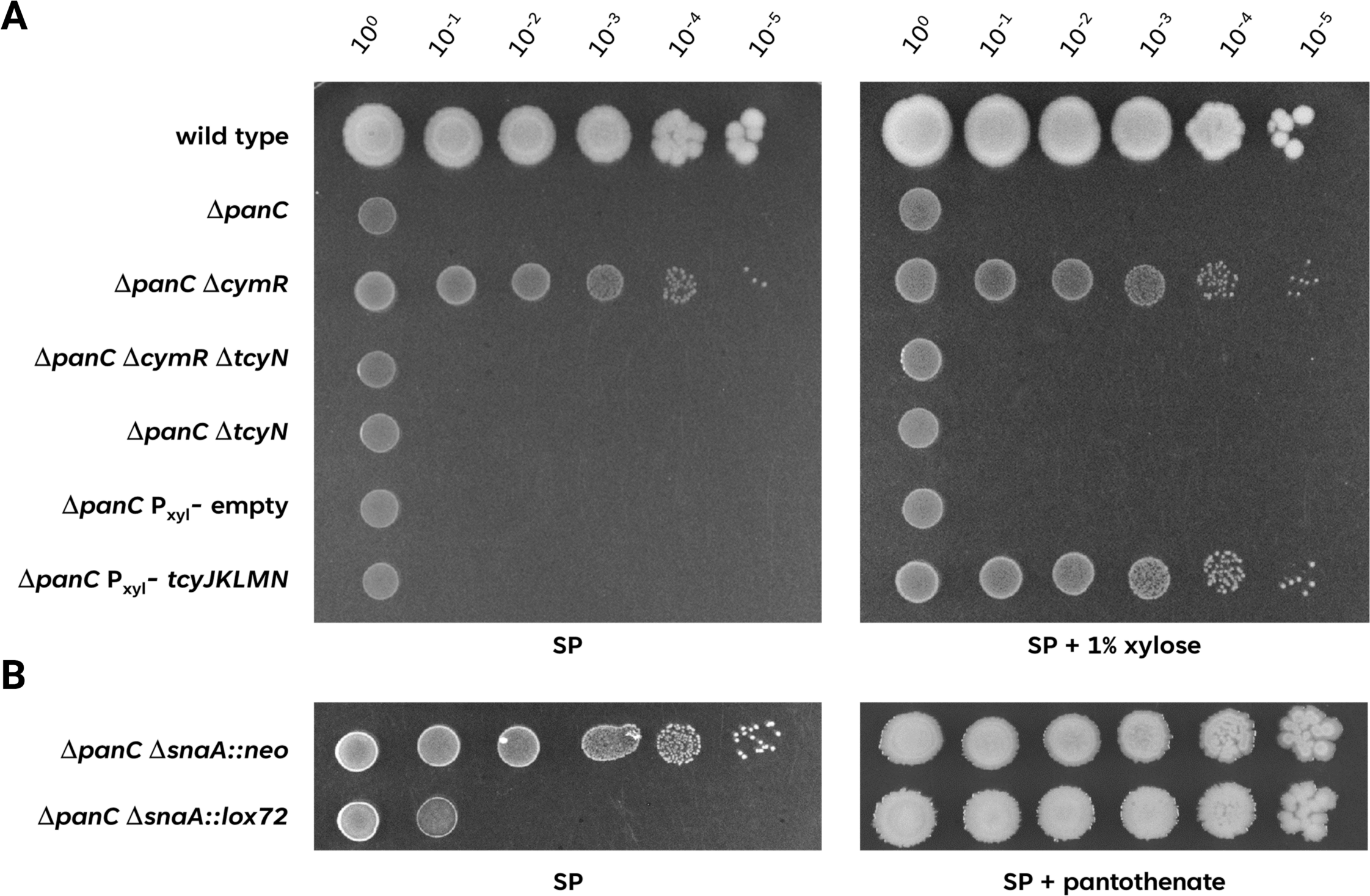
Expression of the TcyJKLMN ABC transporter is responsible and sufficient for the suppression of pantothenate biosynthesis deficiency. A, The growth of the *B. subtilis* wild type was compared with the mutants Δ*panC* (GP4361), Δ*panC* Δ*cymR* (GP4362), Δ*panC* Δ*cymR* Δ*tcyN* (GP4375), Δ*panC* Δ*tcyN* (GP4378), Δ*panC* P*xyl-(empty)* (GP4379) and Δ*panC* P*xyl-tcyJKLMN* (GP4380). The cells were grown in sporulation medium (SP) supplemented with 1 mM pantothenate to an OD_600_ of 1.0, and serial dilutions (10-fold) were prepared. These samples were plated on SP plates with or without the inducer 1% xylose and incubated at 37°C for 48 h. B, The growth of the Δ*panC* Δ*snaA::neo* (GP4364) and the isogenic cre-loxed Δ*panC* Δ*snaA::lox72* (GP4383) was compared. The cells were grown in sporulation medium (SP) supplemented with 1 mM pantothenate to an OD_600_ of 1.0, and serial dilutions (10-fold) were prepared. These samples were plated on SP plates with or without 1 mM of pantothenate and incubated at 37°C for 48 h.

### Modeling of the CysK-CymR repressor complex

Interestingly, CysK does not only act as the cysteine synthase but also as a corepressor of CymR (28). At high intracellular cysteine concentrations, the two proteins form a CymR_2_-CysK_2_ complex (28), and this complex can bind to its DNA target sites and repress transcription. While the crystal structure of the inactive CymR hexamer, which is present in the absence of cysteine, has been solved (PDB 2Y75; 27), no structure is available for the CymR_2_-CysK_2_ repressor complex. We therefore made use of AlphaFold-Multimer (29) to obtain a structural model for the complex (see Fig. 4C). The predicted model exhibited high confidence scores (pLDDT=86.2; pTM=0.771; ipTM=0.669, see http://www.subtiwiki.uni-goettingen.de/v4/predictedComplex?id=170 for an interactive presentation of the complex). The resulting structure suggests that the C-terminus of CymR protrudes like an anchor into CysK. Subsequently, we assessed whether the predicted CymR_2_-CysK_2_ repressor complex would bind to DNA by employing a molecular docking approach with HDOCK (30). Using the dsDNA of the *cysK* upstream region as a template, which contains the CymR binding site (26), we found a strikingly high predicted confidence score for the protein-DNA complex (0.933), suggesting that the CymR_2_-CysK_2_ repressor complex very likely binds to the DNA in this way (see Fig. 4C). Such an interaction would not be possible for the CymR hexamer, as the DNA binding domains are facing inwards (27). We hypothesize that in the absence of cysteine, dimeric apo-CysK interacts with the hexamer, causing its disassembly. As a result, the CymR_2_-CysK_2_ repressor complex is formed, with its DNA binding site in a position to interact with its target DNA (see Extended Data Movie M1).

In the model of the CymR_2_-CysK_2_ repressor complex, the penultimate residue of CymR, Tyr-137 interacts with Gly-223 of CysK. It is tempting to speculate that the truncation of CysK just at this position interferes with the interaction and thus prevents the formation of the CysK_2_-CymR_2_ repressor complex. This idea is indeed supported by the predicted structure of the complex between CymR and the truncated CysK. CymR seems to be completely reorientated with the DNA-binding domain facing towards the truncated CysK which would completely prevent DNA binding (see Extended Data Fig. 3). The absence of CymR-mediated transcription repression resulting from a mutation in the DNA-binding domain or from abortive interaction with the truncated corepressor CysK results in expression of the genes of the CymR regulon (26).

### The cystine transporter TcyJKLMN is required to bypass *panC* essentiality

The results described above suggest that the constitutive expression of the genes and operons of the CymR regulon is responsible for the ability of the *panB* or *panC* suppressor mutants to grow in the absence of added pantothenate. The CymR regulon consists of 42 genes that are organized in 10 transcription units (26, see http://www.subtiwiki.uni-goettingen.de/v4/regulon?id=protein:50930C56C27D22715620A350220E3C56ADB41020, 31). To get an idea which of these genes might be responsible for the suppression, we performed an initial screening by transferring deletion constructs from 25 representative mutants of the Koo mutant collection (12, see Supplementary Table S1) to the *panC cymR* double mutant GP4362. Then we checked the transformants for their ability to grow on plates in the absence of added pantothenate. We expected that the inactivation of genes that are required for the suppression phenotype would result in loss of growth under this condition. While no effect was observed in most cases, the inactivation of the *tcyL*, *tcyM*, and *tcyN* genes reversed the suppression and prevented growth of the *panC cymR* double mutant in the absence of added pantothenate (see Fig. 5A for the *panC cymR tcyN* mutant GP4375). The *tcyJ*, *tcyK*, *tcyL*, *tcyM*, and *tcyN* genes encode subunits of an ABC transporter for the uptake of cystine. To demonstrate that this ABC transporter is indeed responsible for the suppression of the *panC* mutant, we constructed strain GP4380, in which the *tcyJKLMN* genes are expressed under the control of the xylose-inducible promoter of the *B. subtilis xylAB* operon. In this strain, which also lacks the *panC* gene, the addition of the inducer xylose allowed growth in the absence of pantothenate. In contrast, no growth was observed in the isogenic control strain GP4379 that carries the empty plasmid integrated into the genome. Moreover, we tested growth of a *panC* Δ*snaA* mutant from the Koo collection (12). In this library, each reading frame is replaced by an antibiotic cassette with a relatively strong outwardly facing promoter, so overexpression of genes can be the result of the replacement of upstream genes (32). Indeed, the *panC* Δ*snaA* mutant GP4364 was able to grow on complex medium in the absence of added pantothenate (see Fig. 5B). In contrast, the deletion of the resistance cassette including the promoter (GP4383) resulted in loss of growth as the *tcyJKLMN* genes were no longer constitutively expressed (Fig. 5B). Thus, the CymR-independent overexpression of the *tcyJKLMN* genes is sufficient to rescue the *panC* mutant. These results also demonstrate that the mutations affecting CymR and its co-repressor CysK work by allowing constitutive expression of the *tcyJKLMN* genes.

### Identification of the TcyJKLMN substrate that feeds into the coenzyme A salvage pathway

As described above, the TcyJKLMN complex seems to allow growth of *B. subtilis* in the absence of pantothenate biosynthesis. This suggests that this ABC transporter may transport not only cystine, but also a precursor molecule that allows pantothenate-independent synthesis of coenzyme A. This idea is supported by the established promiscuity of many amino acid transporters (16, 33, 34). Candidates for such substrates are pantothenate or pantetheine which could be introduced into the either of the two pathways to CoaA (see above, Fig. 1). Phosphorylated or very large molecules are rather unlikely to be imported by permeases (23), so we focused on pantothenate and pantetheine. To distinguish between the two molecules, we used strains that allowed inducible expression of the *tcyJKLMN* genes in the backgrounds of *panC*, *panC coaA*, or *panC coaX* mutants (see Fig. 6A). As described above (see Fig. 5A), induction of the *tcy* genes by xylose allowed growth of the *panC* mutant (GP4380) on complex medium, suggesting that the transported substrate is present in the medium. In contrast, the isogenic *coaA* mutant GP4444 was unable to grow, whereas the *coaX* mutant strain GP4489 grew like the wild type upon induction of the *tcyJKLMN* genes. This observation indicates that the bifunctional pantothenate/pantetheine kinase CoaA is essential for the growth of the suppressor mutants, whereas the monofunctional CoaX is not sufficient. The addition of pantothenate allowed growth of both the *coaA* and the *coaX* mutant, in good agreement with the retained pantothenate kinase activity in both strains. Pantethine (the oxidized dimeric form of pantetheine), in contrast, rescued growth of the *coaX,* but not of the *coaA* mutant. This result supports the notion that CoaA is required for the suppression of the *panC* mutant by overexpression of the TcyJKLMN ABC transporter. Taken together, we conclude that the suppression occurs via the pantetheine and CoaA-dependent route coenzyme A salvage route (see Fig. 1). This idea is further supported by the observation that both pantothenate and phosphopantothenate can be found in the *B. subtilis* wild type strain, whereas both metabolites are absent from the *panC* mutant (see Fig. 6B). As the *panC* mutant is unable to produce pantothenate irrespective of the expression of the TcyJKLMN ABC transporter, we can exclude pantothenate as a possible substrate for the transporter.

**Figure 6:**
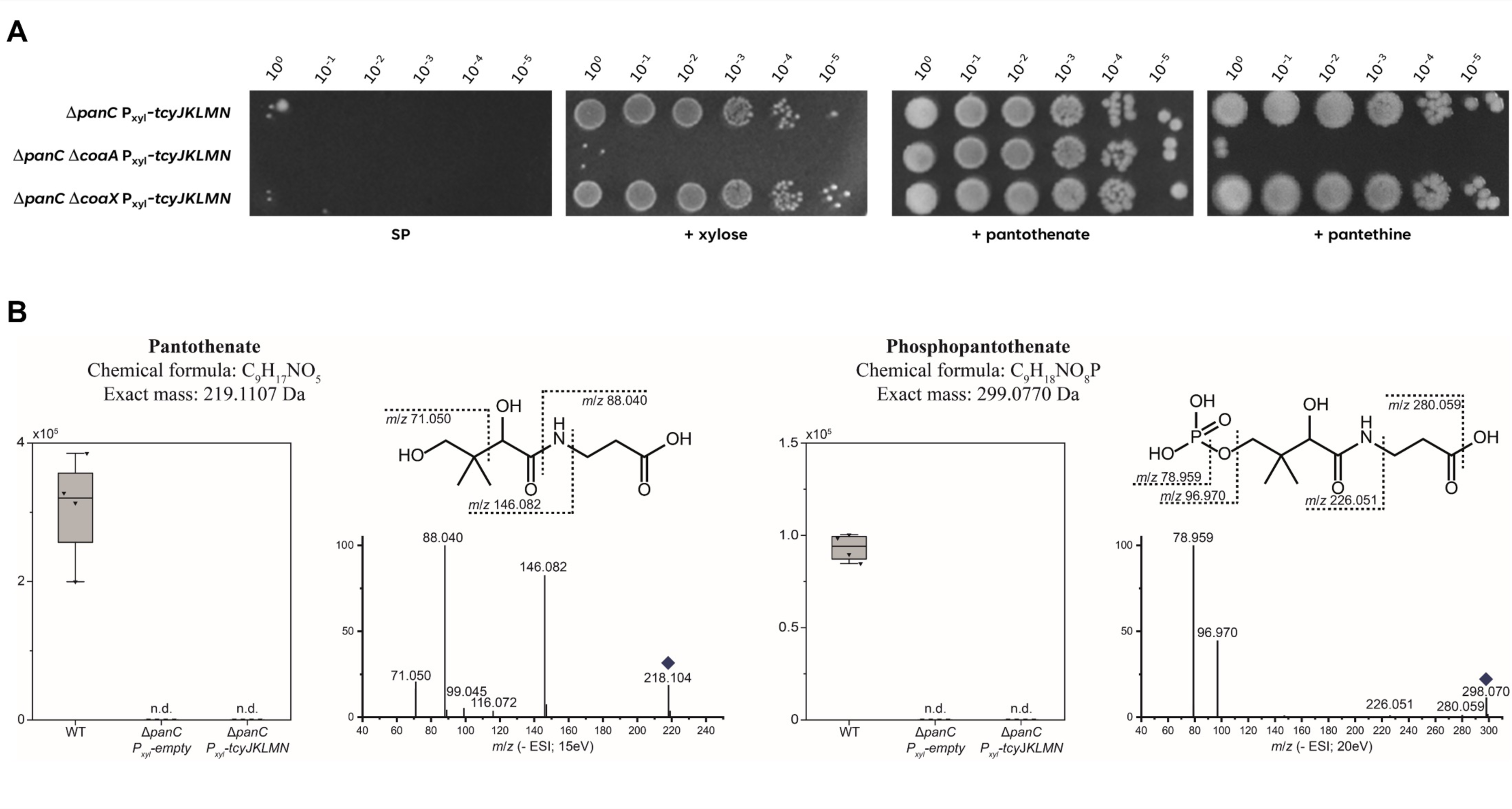
Suppression by expression of the TcyJKLMN ABC transporter depends on CoaA. A, The growth of the Δ*panC* P*xyl-tcyJKLMN* (GP4380) mutant was compared with the isogenic Δ*coaA* (GP4444) and Δ*coaX* (GP4489) mutants. The cells were grown in sporulation medium (SP) supplemented with 1 mM pantothenate to an OD_600_ of 1.0, and serial dilutions (10-fold) were prepared. These samples were plated on SP plates with or without the inducer 1% xylose or supplemented with 250 µM pantothenate or pantethine and incubated at 37°C for 48 h. B, *B. subtilis* extracts of WT, *panC P_xyl_-empty* (GP4379) and *panC P_xyl_-tcyJKLMN* (GP4380) were analyzed by ultrahigh performance liquid chromatography coupled to high resolution mass spectrometry. Relative intensities of four independent experiments are shown as boxplots. Pantothenate and phosphopantothenate were detected in the WT extracts but not in the *panC* mutant strains (n.d.). The identity of either pantothenate or phosphopantothenate in the extracts was confirmed by MS/MS analyses and authentic standard for pantothenate. Respective fragmentation spectra at 15 eV or 20 eV are shown for negative ionization mode. The precursor ion is marked by a rhomb. Characteristic fragments are displayed in the respective structures.

The exclusion of pantothenate suggests that instead pantetheine (or its oxidation product pantethine) may be transported by the ABC transporter. To get support for this idea, we analyzed the *panC* mutant GP4361, the isogenic strain GP4364 overexpressing the *tcyJKLMN* genes due to the replacement of the *snaA* gene, and the *panC* mutant also lacking the *tcyJKLMN* genes (GP4660). Due to the essentiality of *panC* and the lack of any possible substrate for uptake by the TcyJKLMN ABC transporter, these strains are unable to grow on minimal medium in the absence of any precursor molecules for coenzyme A. We then tested the growth of the strains at different concentrations of pantetheine and pantethine (see Extended Data Fig. 4). All strains were able to grow at high concentrations of pantetheine and pantethine (50 µM). However, the ability to grow diminished with decreasing pantetheine and pantethine concentrations and no growth was possible at concentrations below 2.5 µM and 5 µM for pantetheine and pantethine, respectively, independent of the presence or absence of the TcyJKLMN ABC transporter (see Extended Data Fig. 4). Thus, we had to exclude the possibility that pantetheine/ pantethine might be transported by TcyJKLMN.

As the two obvious hypotheses had been discarded by the experimental data, we had to consider alternative possibilities. The TcyJKLMN transporter is a high-affinity cystine transporter (35). Cystine is formed by oxidation of two cysteines yielding the homodimeric disulfide. Similarly, pantethine is the product of pantetheine oxidation which consists of two pantetheine molecules linked by an S-S bridge. Taking into account that the TcyJKLMN transporter probably recognizes a cysteine-moiety with a disulfide bond and that pantetheine is required for CoaA-mediated coenzyme A salvage, we hypothesized the presence of a heterodimeric disulfide formed by cysteine and pantotheine under oxidizing conditions (see Extended Data Fig. 5). Such a molecule might then be taken up by TcyJKLMN and subsequently be reduced in the cell to cysteine and pantetheine.

To test the spontaneous formation of a disulfide hybrid molecule of cysteine and pantetheine, we mixed both molecules with a 50-fold excess of cysteine and incubated the mixture overnight. Similarly, we incubated both cysteine and pantetheine alone. The analysis of the resulting products by UHPLC-HRMS/MS (see Fig. 7A, Extended Data Fig. 6), showed the expected oxidation products cystine and pantethine were found upon the incubation of cysteine or pantetheine, respectively. These products were also detected after the incubation of the mixture of cysteine and pantetheine. In addition, and as suspected, we identified the heterodimeric cysteinopantetheine.

**Figure 7:**
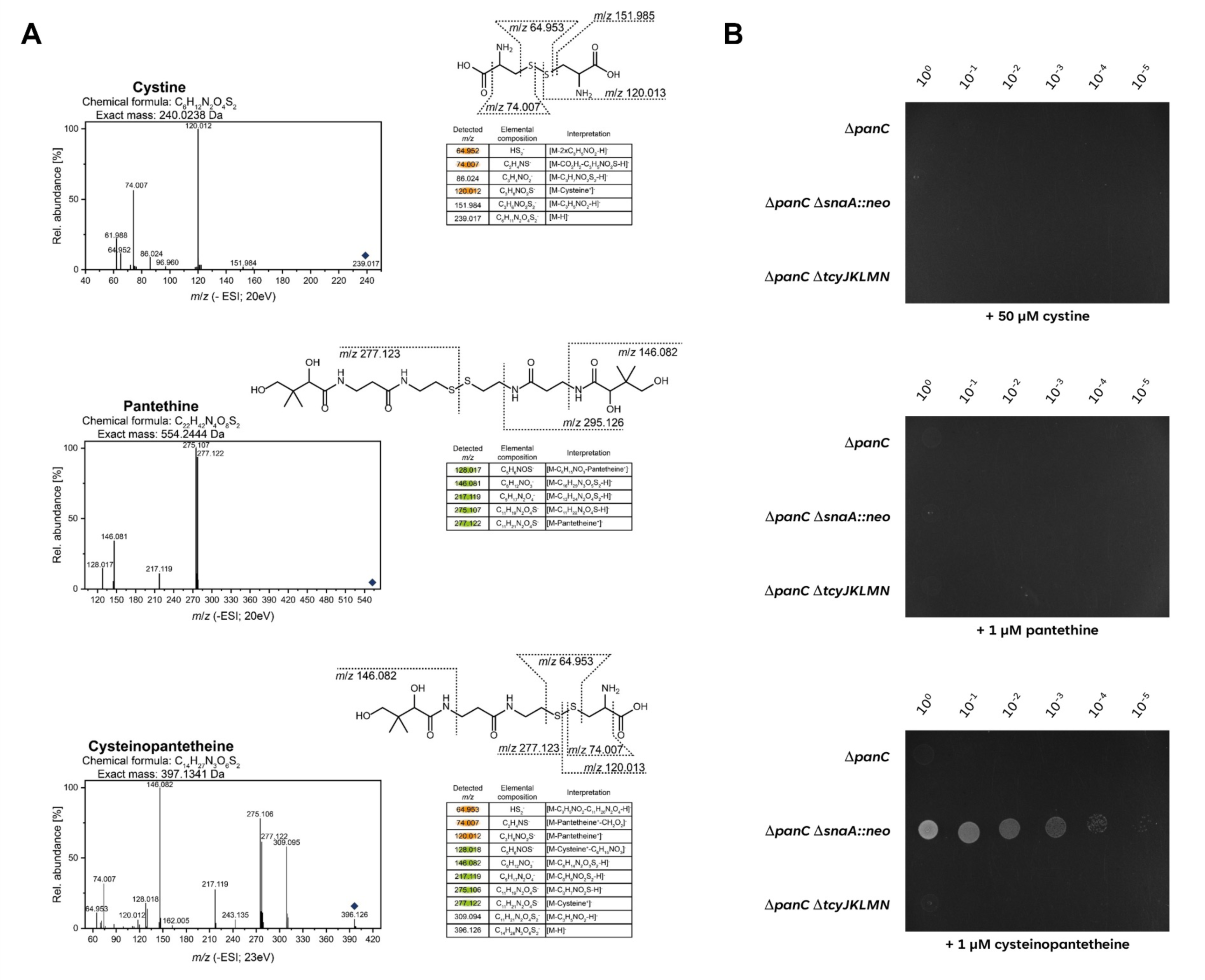
The ABC Transporter TcyJKLMN is active in the transport of cysteine-pantetheine. A, The identity of cysteinopantetheine was confirmed by comparative MS/MS analyses with cystine and pantethine. Authentic standards for (top panel) cystine and (middle panel) pantethine as well as (bottom panel) a solution containing cysteinopantetheine were analyzed by ultra high performance liquid chromatography coupled to high resolution mass spectrometry and fragmented by collision induced dissociation at 20 eV (A, B) or 23 eV (C) in negative ionization mode. The respective MS/MS spectra were interpreted as shown in the tables and as characteristic fragments in the structure. A cysteine moiety is considered as C_3_H_6_NO_2_S and a pantetheine moiety is considered as C_11_H_21_N_2_O_8_S. The respective precursor ions are marked by a rhomb. Characteristic fragments used for structure confirmation of cysteinopantetheine are marked in orange and green, respectively. B, The growth of the Δ*panC* (GP4361) mutant was compared with the isogenic Δ*panC* Δ*snaA* (GP4364) and Δ*panC* Δ*tcyJKLMN* (GP4660). The cells were grown in C Glc minimal medium supplemented with 1 mM pantothenate to an OD_600_ of 1.0, and serial dilutions (10-fold) were prepared. These samples were plated on C Glc minimal medium plates 50 µM oxidized cysteine (top), 1 µM oxidized pantetheine mix (middle), or 1 µM of oxidized cysteine/pantetheine mix containing cysteine-pantetheine (bottom) and incubated at 37°C for 48 h.

Next, we tested the growth of the three *panC* mutants with standard or high-level expression of *tcyJKLMN* or in the absence of the ABC transporter on minimal medium supplemented with the final oxidation products of cysteine (corresponding to 50 µM cysteine), pantetheine (corresponding to 1 µM pantetheine), and the mixture of both molecules (the same concentrations for both metabolites). As shown in Fig. 7B, no growth was possible with the oxidation products of cysteine (cystine) or pantetheine (pantethine) alone. This is not unexpected, as we already have hypothesized that a hybrid molecule that sufficiently resembles cysteine for transport but that also brings pantetheine is required for the TcyJKLMN-dependent bypass of the growth defect of the *panC* mutant. Indeed, the addition of the oxidation product of the cysteine/ pantetheine mixture that contained the novel molecule cysteinopantetheine allowed growth of the *panC* mutant overexpressing the TcyJKLMN ABC transporter. We can therefore assume that the complex medium contains small amounts of pantetheine as well as cysteine, and that these molecules can spontaneously form cysteinopantetheine which is then taken up by TcyJKLMN. Upon reduction in the cytoplasm, the pantetheine then seems to feed into the CoaA-dependent coenzyme A salvage (see Fig. 1).

## Discussion

With their involvement in more than 30 reactions, among them in the intersection between glycolysis and the citric acid cycle and the initial reaction of fatty acid biosynthesis, coenzyme A and its derivative acetyl-CoA are central to the physiology of *B. subtilis* (see https://corewiki.uni-goettingen.de/metabolite/4547 and https://corewiki.uni-goettingen.de/metabolite/4710). It is therefore not surprising that *B. subtilis* has developed two independent routes for the acquisition of coenzyme A, and that the bacterium encodes transporters for multiple precursor molecules to import rather than to synthesize them (see Fig. 1).

In principle, *B. subtilis* can synthesize coenzyme A from pantothenate or from pantetheine. While pantothenate can be made *de novo* from intermediates of central carbon metabolism such as pyruvate and oxaloacetate (via aspartate), there is no pathway for the biosynthesis of pantetheine. Instead, pantetheine is a degradation product of the phosphopantetheine moiety of the acyl carrier protein AcpA which plays a major role in fatty acid biosynthesis (36). Thus, the pantetheine-dependent salvage pathway to coenzyme A is only possible if pantetheine or a source of it can be taken up.

In this work, we have identified all the players in the biosynthesis of coenzyme A in *B. subtilis*. Interestingly, two steps in the pantothenate-dependent pathway can be catalyzed by more than one enzyme. Three enzymes can reduce ketopantoate to pantoate, the ketopantoate reductases PanE and PanG as well the ketol-acid reductase IlvC which is also part of branched-chain amino acid biosynthesis. Many organisms have multiple ketopantoate reductases, however, it is so far unknown whether these enzymes have distinct activities. In many bacteria that contain only one of these enzymes, such as *E. coli*, *Listeria monocytogenes*, *Bacillus licheniformis* or *Streptococcus pyogenes*, only PanE is present suggesting that this is the major player in pantoate production. This idea is supported by the observation that the *panE* gene of *B. subtilis* is more strongly expressed than *panG* under most conditions (37). Two enzymes, CoaA and CoaX, can phosphorylate pantothenate. Unlike the ketopantoate reductases which are member of the same cluster of orthologous genes (38), CoaA and CoaX are members of different protein families. Most bacteria contain only one of these enzymes which is typically CoaX, as in cyanobacteria, clostridia, bacteroidetes, or the spirochaetes. In gamma-proteobacteria, most species have either CoaA or CoaX. Only in the bacilli and actinobacteria, many species have both enzymes. Interestingly, the lactic acid bacteria encode CoaA rather than CoaX, whereas bacteria of the genus *Staphylococcus* have neither CoaA nor CoaX, but a third type, CoaW (39). The two enzymes of *B. subtilis* are distinct from each other as CoaA and other enzymes of this family can phosphorylate multiple pantothenate analogues, including pantetheine whereas member of the CoaX-like family are highly selective for pantothenate (23, 40, 41).

For metabolites with complex biosynthetic pathways, bacteria often use salvage pathways to recycle broken or semi-complete precursor molecules in addition to the more costly complete *de novo* synthesis. In *B. subtilis*, this is the case for purine nucleotides, for the sulphur-containing amino acids methionine and cysteine (42, 43, 44) and also for coenzyme A as shown in this work. This requires the efficient uptake of these incomplete precursors which are often available in media as a result of the death of other cells and the concomitant release of metabolites. However, metabolites that have a low abundance in cells will then also be present at very low concentrations in media, suggesting the requirement for high-affinity uptake systems. In this study, we have identified uptake systems for pantothenate and cysteinopantetheine.

Pantothenate is transported by an ECF transporter that obtains energy derived from ATP hydrolysis by the two cytoplasmic subunits EcfA1 and EcfA2. The membrane spanning transporter EcfT interacts both with the ATPase subunits of the complex and with the substrate-specific PanU protein (see Fig. 3C). The ECF transporters are required for the high-affinity uptake of vitamins and cofactors as well as of other rare nutrients such as tryptophan or cobalt. They are typically composed of the general EcfA1, EcfA2, and EcfT proteins, and substrate-specific binding S proteins (21). *B. subtilis* contains six S proteins, as judged from sequence similarity and functional analyses. While the substrates for four of these proteins have been identified prior to this work, no functional analyses had been performed for PanU (YhfU) and YuiG. Interestingly, these two proteins share 26.8% amino acid identity, and both are subject to repression of their genes by the bifunctional biotin-protein ligase and transcription factor BirA (45). Indeed, the proteins are similar to biotin-specific S proteins suggesting a role in biotin uptake. Most bacteria that contain proteins of this family, encode only one of them. Interestingly, several bacteria of the Bacilli group contain two, and in some cases even three members of this family. It is tempting to speculate that these proteins have diverged in evolution to serve the transport of different substrates – biotin and pantothenate. It will thus be interesting to test the substrate specificity of YuiG, the PanU paralog in *B. subtilis*.

Our suppressor screen of mutants defective in pantothenate synthesis yielded mutations affecting the expression of an ABC transporter previously implicated in the uptake of cystine (35). Since coenzyme A is essential for bacterial growth, these mutations already suggested the possibility of transport of a precursor of coenzyme A by the TcyJKLMN ABC transporter. High expression of this transporter allowed the uptake of this unknown intermediate from complex medium. However, the transporter was not implicated in the uptake of the potential precursor molecules pantothenate, pantetheine, and pantethine. Moreover, more than 5 µM pantetheine or even more than 50 µM of pantetheine are required for growth in the absence of pantothenate biosynthesis (see Extended Data Fig. 4). The presence of a precursor molecule in complex medium which can be taken up by TcyJKLMN suggested that the ABC transporter accumulated a different molecule from the medium. It is reasonable to assume that the concentration of pantetheine in complex medium is very low whereas amino acids are likely to be much more abundant. Thus, we considered the possibility that under oxidizing conditions a hybrid molecule between cysteine and pantetheine could be formed and then be taken up by the TcyJKLMN transporter. As many amino acid transporters are not very substrate-specific and cystine and cysteinopantetheine even share the cysteine moiety of the molecule, this idea seemed highly attractive, and could be proven by experimental analysis. Cysteinopantetheine is a novel metabolite that has not been described so far.

It is tempting to speculate that more molecules might form hybrids under specific conditions, and that the growth requirement for one moiety of such hybrids and the possibility to transport it for the second one, may facilitate a rich underground metabolism which still awaits its exploration.

## Materials and methods

### Bacterial strains, growth conditions, and phenotypic characterization

E. coli DH5α (46) was used for cloning. All *B. subtilis* strains used in this study are listed in Table S1. All strains are derived from the laboratory strain 168 (*trpC2*). *B. subtilis* and *E. coli* was grown in Lysogeny Broth (LB medium) in sporulation medium (SP) (46, 47). For testing the requirement for pathway intermediates, C minimal medium supplemented with 0.5% glucose (C Glc) (48) was used. LB, SP or C Glc plates were prepared by addition of 17 g Bacto agar/l (Difco) to LB and SP, respectively.

### DNA manipulation and genome sequencing

*B. subtilis* was transformed with plasmids, genomic DNA or PCR products according to the two-step protocol (46, 47). Transformants were selected on SP plates containing erythromycin (2 µg/ml) plus lincomycin (25 µg/ml), tetracycline (12.5 µg/ml), chloramphenicol (5 µg/ml), kanamycin (10 µg/ml), or spectinomycin (150 µg/ml). Phusion DNA polymerase (Thermo Fisher Scientific, Dreieich, Germany) was used as recommended by the manufacturer. DNA fragments were purified using the peqGOLD Cycle-Pure Kit (Peqlab, Erlangen, Germany). DNA sequences were determined by the dideoxy chain termination method (46). Chromosomal DNA from *B. subtilis* was isolated using the peqGOLD Bacterial DNA Kit (Peqlab, Erlangen, Germany). To identify the mutations in the suppressor mutants, their genomic DNA was subjected to whole-genome sequencing. Briefly, the reads were mapped on the reference genome of *B. subtilis* 168 (GenBank accession number: NC_000964) (49). Mapping of the reads was performed using the Geneious software package (Biomatters Ltd., New Zealand) (50). Frequently occurring hitchhiker mutations (51) and silent mutations were omitted from the screen. The resulting genome sequences were compared to that of our in-house wild type strain. Single nucleotide polymorphisms were considered as significant when the total coverage depth exceeded 25 reads with a variant frequency of ≥90%. All identified mutations were verified by PCR amplification and Sanger sequencing.

### Plasmids

pDR244 was used to delete the resistance cassettes in the strains of the Koo collection as described. The deletion results in a cre-lox scar (12). The integrative plasmid pGP886 (52) for the xylose-inducible expression of genes in *B. subtilis*. Upon transformation, the plasmid integrates into the *xkdE* gene of prophage PBSX. To express the tcyJKLMN genes under the control of the xylose-inducible promoter, we constructed plasmid pGP4018 as follows: the *tcyJKLMN* genes were amplified using appropriate primers. The DNA fragment was digested with XbaI and EcoRI (introduced with the PCR primers) and ligated into pGP886 linearized with the same enzymes.

### Construction of mutant strains by allelic replacement

Deletion of the *coaBC*, *panC,* and *tcyJKLMN* genes was achieved by transformation of *B. subtilis* 168 with PCR products constructed using oligonucleotides to amplify DNA fragments flanking the target genes and an appropriate intervening resistance cassette as described previously (53). The integrity of the regions flanking the integrated resistance cassette was verified by sequencing PCR products of about 1,100 bp amplified from chromosomal DNA of the resulting mutant strains (see Table S1).

### UHPLC-HRMS analysis of *B. subtilis* extracts and cysteinopantetheine

*B. subtilis* was grown to midlogarithmic phase and 15 OD units harvested by centrifugation. Cells were washed once with Tris-HCl (10 mM, pH7.0) and then resolved in methanol:acetonitrile:water (40:40:20 (*v/v/v*)). Cells were further disrupted by ultrasonic bath for 30 min. Cell debris was removed by centrifugation at 18.000 x g at room temperature. The supernatant was transferred into micro vials and analyzed by ultrahigh performance liquid chromatography (UHPLC, 1290 Infinity, Agilent Technologies) coupled to a high-resolution mass spectrometer (HRMS, 6540 or 6546 UHD Accurate-Mass Q-TOF LC-MS instrument with Dual Jet Stream Technology, Agilent Technologies) as described (54). The identity of pantothenate was confirmed by an authentic standard as well as the comparison of the MS/MS spectra to those of the NIST Mass Spectral Library (https://chemdata.nist.gov/), NIST#: 1351032 (positive ionization), NIST#: 1351077 (negative ionization). Phosphopantothenate was identified by accurate mass analysis and interpretation of HRMS/MS spectra in negative ionization with *m/z* 78.959 and *m/z* 96.970 as analytical fragments of the phosphate moiety. The detection limit of pantothenate was determined to be 10 fmol per injection with an authentic standard.

The structure of cysteinopantetheine was confirmed by comparative MS/MS analyses under consideration of the characteristic fragments obtained from MS/MS spectra of cystine and pantethine. These reference spectra were obtained from MS/MS fragmentation of authentic standards and were further compared to high resolution MS/MS spectra from the NIST Mass Spectral Library for cystine, NIST#: 1190075 (negative ionization) and NIST#: 1190045 (positive ionization); for pantethine, NIST#: 1271189 (positive ionization). Based on the characteristic fragments of both homodimeric disulfides the identity of cysteinopantetheine was confirmed.

### Visualization of global mutant fitness data

For all non-essential genes of *B. subtilis*, the fitness of the corresponding mutants as compared to a wild type strain has been determined under several growth conditions in complex and minimal media (12). To make this information accessible in an intuitive way, we added a “Relative mutant fitness” widget to the gene pages of the database *Subti*Wiki (31). The fitness scores for each gene and condition were extracted and stored in the *Subti*Wiki database. For the visualization, a bar plot provided by the Apache ECharts library (v4.1.0) was used.

### Atomic model generation with ColabFold

The local installation of ColabFold (55) (version 1.5.5, released on January 30, 2024), which is compatible with AlphaFold 2.3.2, was used to predict atomic models of the CymR_2_-CysK_2_ heterotetramer and the EcfA1/ EcfA2/ EcfT/ PanU complex. For the jobs, the default options were kept. A total of 5 unrelaxed atomic models of CymR_2_-CysK_2_ and EcfA1/A2/T/PanU were calculated and subsequently analyzed. The models were evaluated using the PAE Viewer webserver (56). The models were ranked from 1 to 5 according to Alphafold-Multimer’s model confidence metric, a weighted combination of ipTM and pTM. The predicted template modeling score (pTM) measures the structural congruence between two folded protein structures, while the ipTM (interface pTM) scores interactions between residues of different chain to estimate the accuracy of interfaces. The pLDDT (predicted local-distance difference test) is a confidence measure for the per-residue accuracy of the structure.

The predicted model of the CymR_2_-CysK_2_ repressor complex was used as a template for a molecular docking with HDOCK (30) using dsDNA of the CymR binding site of the *cysK* gene (5’-CATAATACCAATACAAATAGTCGGAAATTGAGGT). For the movie, PyMOL 2.5 (http://www.pymol.org/) was used with the models of the CymR_2_-CysK_2_ repressor complex with and without DNA and a crystal structure of the hexameric CymR (PDB: 2Y75). A morphing animation was created between the C and D chains of the CymR hexamer and the CymR dimer of the CymR_2_-CysK_2_ repressor complex model.

## Supporting information

Table S1: Strains used in this study

## Acknowledgements

We are grateful to Constantin Büttner for the help with some experiments. This work was supported by the Deutsche Forschungsgemeinschaft via the Priority Program SPP1879 (project Stu 214/16-2) and Collaborative Research Center CRC1565 (project 469281184) to J.S.. I.F. and M.K. were supported by the Deutsche Forschungsgemeinschaft (GRK 2172 PRoTECT, INST 186/822-1 and INST 186/1434-1).

## Author contributions

RW, MK, and JS conceived and designed the experiments. RW, MK, CH, JD, and KF performed the experiments. RW, MK, CH, CE, JD, KF, IF and JS analyzed and discussed the data; RW, MK, CE, KF, IF and JS wrote the manuscript. All authors edit and approved the manuscript.

## FIGURE LEGENDS

**Box 1.**
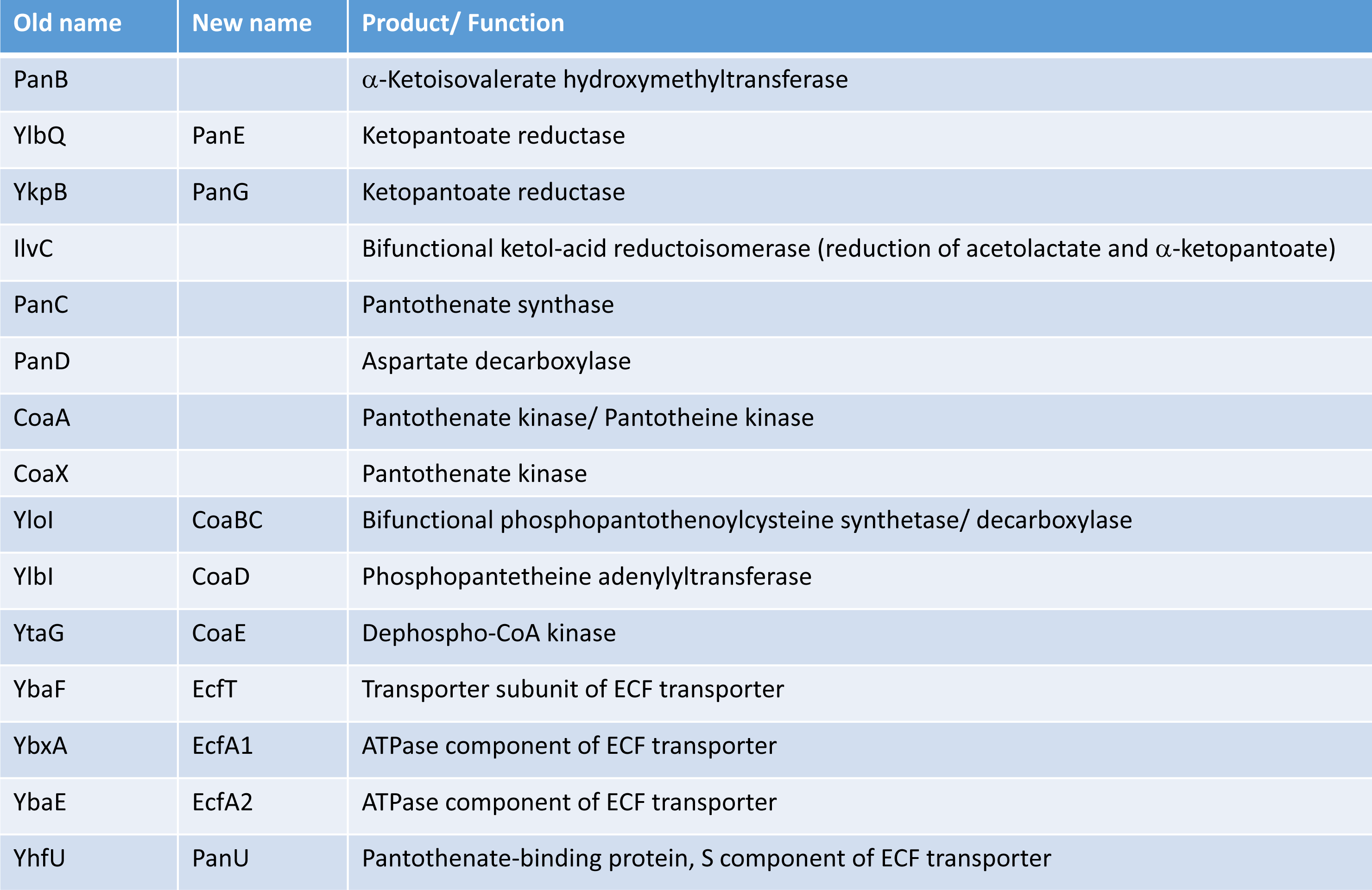
Enzymes involved in the coenzyme A biosynthesis and in the uptake of pantothenate. Enzymes identified in this study were renamed as indicated.

**Extended Data Figure 1:**
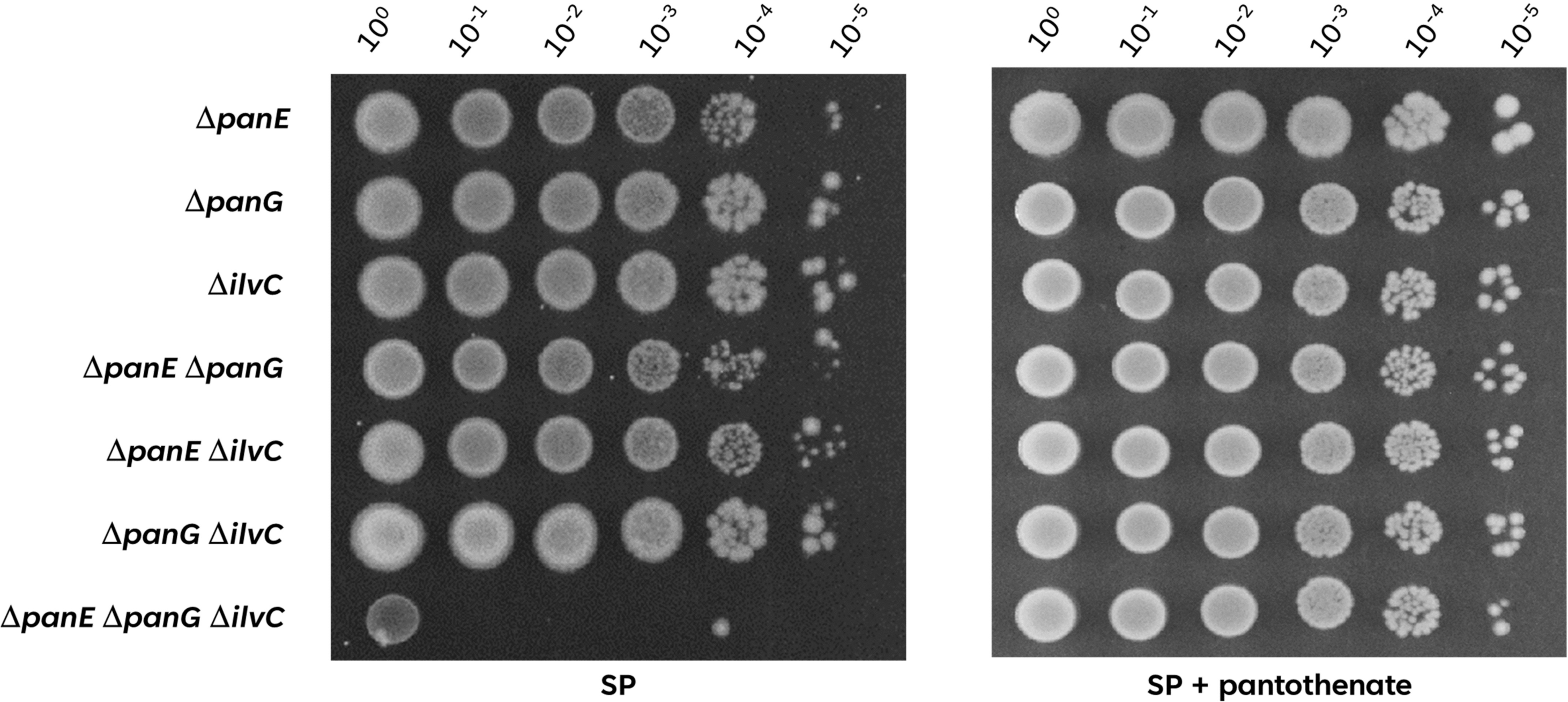
The three enzymes PanE, PanG and IlvC are involved in pantoate biosynthesis. The growth of the *B. subtilis* deletion mutants Δ*panE* (GP3384), Δ*panG* (GP3383), Δ*ilvC* (GP4404), Δ*panE* Δ*panG* (GP3343), Δ*panE* Δ*ilvC* (GP3397), Δ*panG* Δ*ilvC* (GP3346) and Δ*panE* Δ*panG* Δ*ilvC* (GP4403) was compared. The cells were grown in sporulation medium (SP) supplemented with 1 mM pantothenate to an OD_600_ of 1.0, and serial dilutions (10-fold) were prepared. These samples were plated on SP with or without added 1 mM pantothenate and incubated at 37°C for 48 h.

**Extended Data Figure 2:**
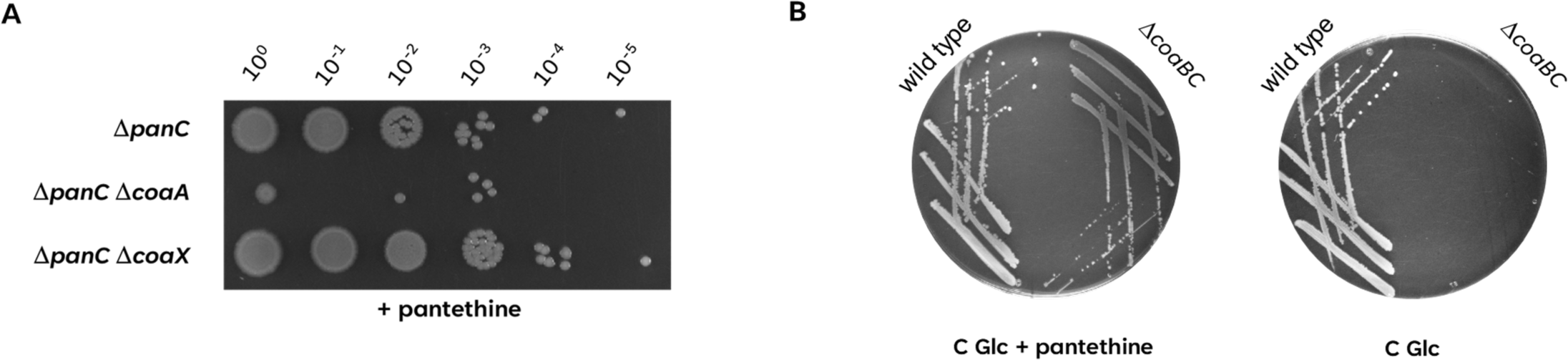
*B. subtilis* harbors a CoaA- and pantetheine-dependent *coaBC* bypass of the biosynthetic pathway. A, The growth of the Δ*panC* mutant (GP4379) was compared with the isogenic Δ*panC* Δ*coaA* (GP4652) and Δ*panC* Δ*coaX* (GP4653) mutants. The cells were grown in sporulation medium (SP) supplemented with 1 mM pantothenate to an OD_600_ of 1.0, and serial dilutions (10-fold) were prepared. These samples were plated on C Glc minimal medium supplemented with 250 µM pantethine and incubated at 37°C for 48 h. B, The deletion mutant Δ*coaBC* GP4670 was streaked out on C Glc minimal medium plates with or without added 250 µM pantethine. The plates were incubated at 37°C for 24h.

**Extended Data Figure 3:**
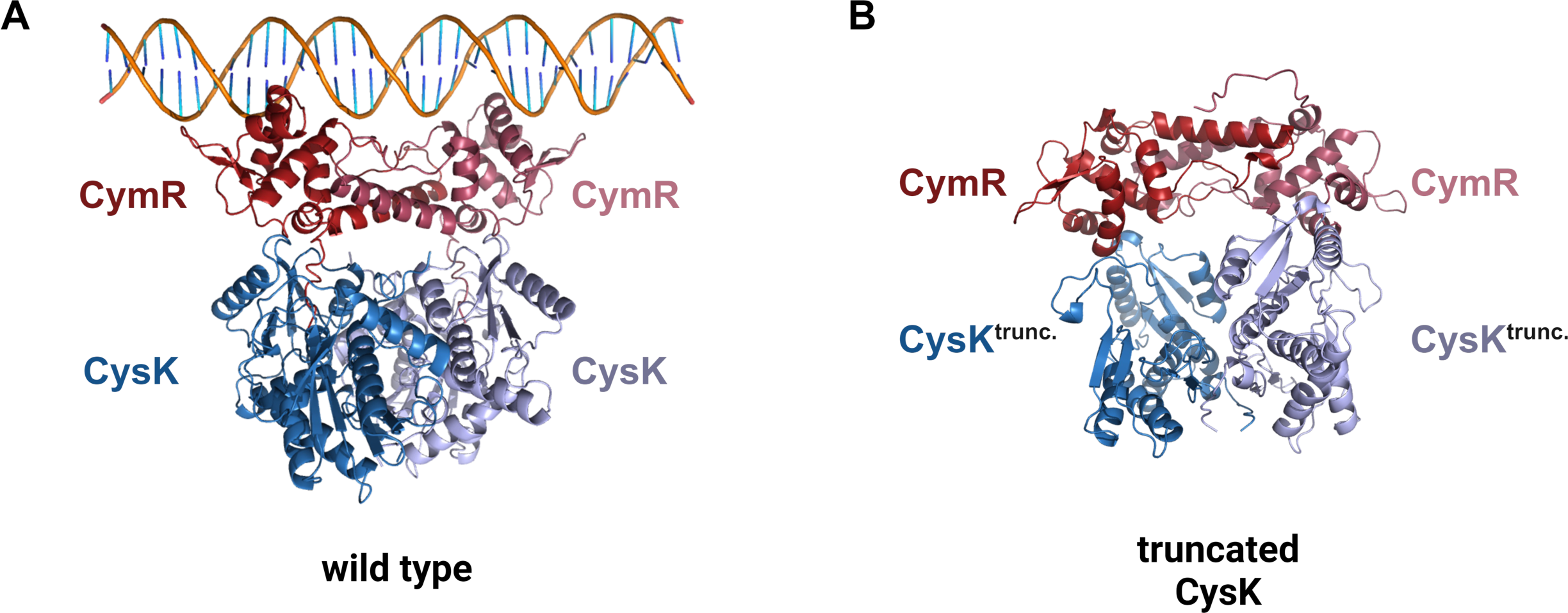
Effect of the CysK truncation on the interaction with CymR. ColabFold/ AlphaFold predicted model of Cym dimer (red, purple) in complex with the full length (A) and truncated (B) CysK dimer (light blue, dark blue). For the wild type model (A), the 5’ UTR of the *cysK* gene was docked with HDOCK.

**Extended Data Figure 4:**
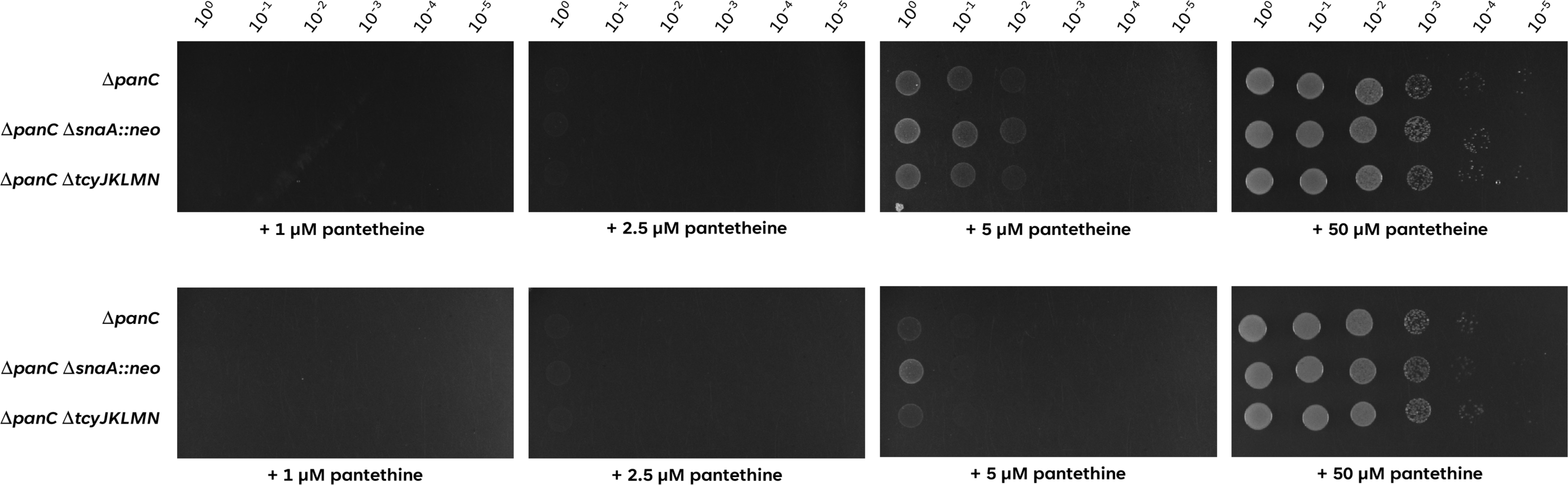
TcyJKLMN is not active in the transport of pantetheine or pantethine. The growth of the Δ*panC* mutant (GP4361) was compared with the isogenic Δ*panC* Δ*snaA::neo* (GP4364) and Δ*panC* Δ*tcyJKLMN* (GP4660), overexpressing or lacking the ABC transporter TcyJKLMN respectively. The cells were grown in C Glc minimal medium supplemented with 1 mM pantothenate to an OD_600_ of 1.0, and serial dilutions (10-fold) were prepared. These samples were plated on C Glc minimal medium plates supplemented with 1 µM, 2.5 µM, 5 µM or 50 µM of either pantetheine or pantethine. The plates were incubated at 37°C for 24h.

**Extended Data Figure 5:**
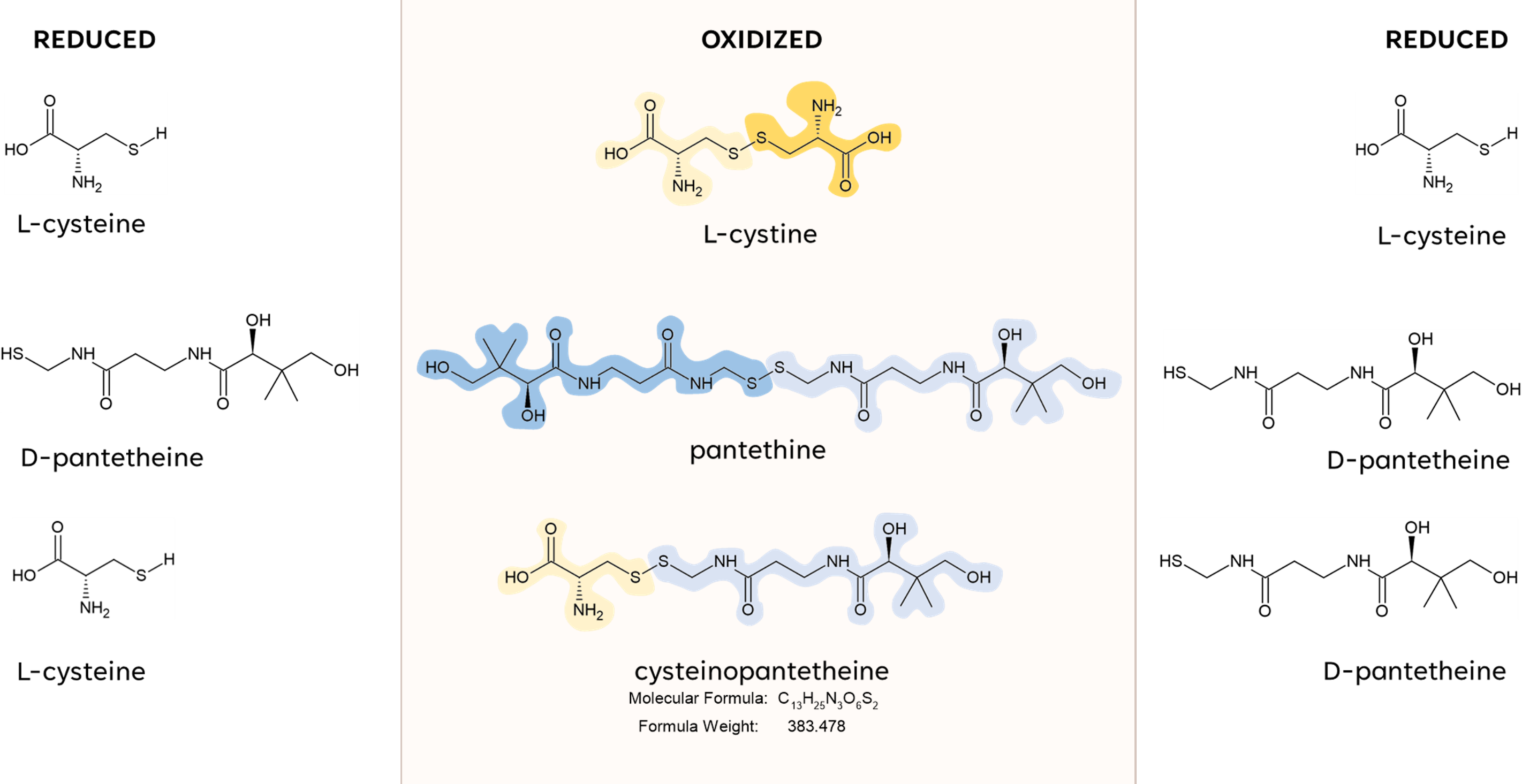
Molecular Structures of the reduced monomers L-cysteine and D-pantetheine and their oxidized dimeric forms L-cystine, pantethine and cysteinopantetheine. The moieties are colorized to enhance the visibility of the dimeric form, for cysteine in dark/light yellow and pantetheine in dark/light blue.

**Extended Data Fig. 6:**
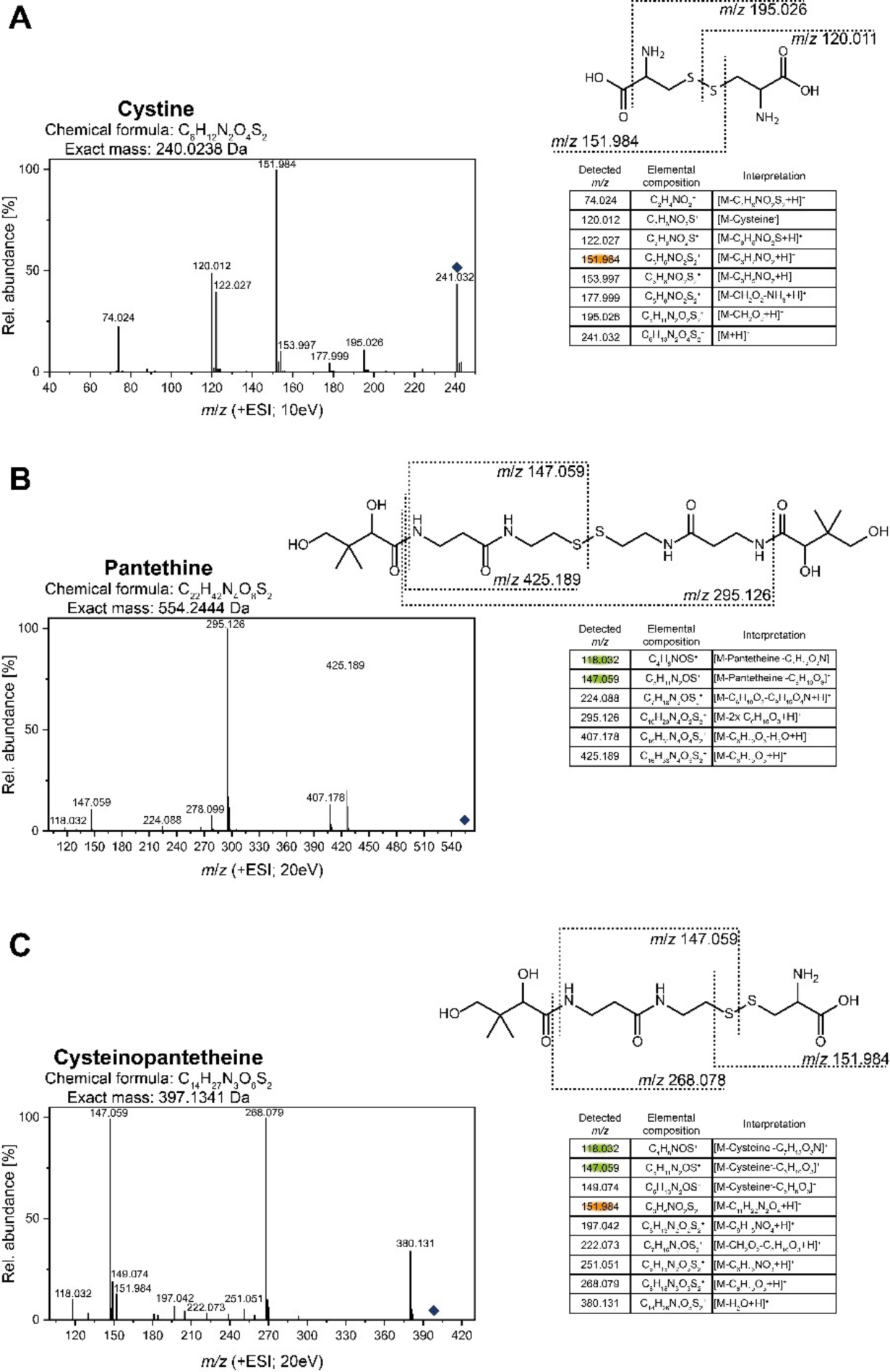
The identity of cysteinopantetheine was confirmed by comparative MS/MS analyses with cystine and pantethine. Authentic standards for (A) cystine and (B) pantethine as well as (C) a solution containing cysteinopantetheine were analyzed by ultra-high performance liquid chromatography coupled to high resolution mass spectrometry and fragmented by collision induced dissociation at 10 eV (A) or 20 eV (B,C) in positive ionization mode. The respective MS/MS spectra were interpreted as shown in the tables and as characteristic fragments in the structure. A cysteine moiety is considered as C_3_H_6_NO_2_S and a pantetheine moiety is considered as C_11_H_21_N_2_O_8_S. The respective precursor ions are marked by a rhomb. Characteristic fragments used for structure confirmation of cysteinopantetheine are marked in orange and green, respectively.

