## Supplementary material for "Coenzyme A biosynthesis in *Bacillus subtilis*: Discovery of a novel precursor metabolite for salvage and its uptake system": Table S1: Strains used in this study

| Strain | Genotype | Reference |
|---|---|---|
| 168 | *trpC2* | Laboratory collection |
| BKK00700 | *trpC2 ΔcoaX::neo* | 12 |
| BKK00730 | *trpC2 ΔcysK::neo* | 12 |
| BKK01470 | *trpC2 ΔecfT::neo* | 12 |
| BKK04520 | *trpC2 ΔydbM::neo* [1] | 12 |
| BKK07190 | *trpC2 ΔyezD::neo* [1] | 12 |
| BKK07200 | *trpC2 ΔyetJ::neo* [1] | 12 |
| BKK09130 | *trpC2 ΔtcyP::neo* [1] | 12 |
| BKK10010 | *trpC2 ΔtrpP::neo* | 12 |
| BKK10370 | *trpC2 ΔpanU:neo* | 12 |
| BKK14440 | *trpC2 ΔpanG::neo* | 12 |
| BKK15110 | *trpC2 ΔpanE::neo* | 12 |
| BKK15580 | *trpC2 ΔcysP::neo* [1] | 12 |
| BKK15610 | *trpC2 ΔylnD::neo* [1] | 12 |
| BKK15620 | *trpC2 ΔsirB::neo* [1] | 12 |
| BKK15630 | *trpC2 ΔylnF::neo* [1] | 12 |
| BKK21980 | *trpC2 ΔypdP::neo* | 12 |
| BKK22410 | *trpC2 ΔpanD::neo* | 12 |

| | | |
|---|---|---|
| BKK22420 | *trpC2 ΔpanC::neo* | 12 |
| BKK22430 | *trpC2 ΔpanB::neo* | 12 |
| BKK23050 | *trpC2 ΔribU::neo* | 12 |
| BKK23760 | *trpC2 ΔcoaA::neo* | 12 |
| BKK27240 | *trpC2 ΔyrhC::neo* [1] | 12 |
| BKK27260 | *trpC2 ΔmccA::neo* [1] | 12 |
| BKK27270 | *trpC2 ΔmtnN::neo* [1] | 12 |
| BKK27280 | *trpC2 ΔyrrT::neo* [1] | 12 |
| BKK27520 | *trpC2 ΔcymR::neo* | 12 |
| BKK28290 | *trpC2 ΔilvC::neo* | 12 |
| BKK29280 | *trpC2 ΔytnM::neo* [1] | 12 |
| BKK29290 | *trpC2 ΔsndA::neo* [1] | 12 |
| BKK29300 | *trpC2 ΔribR::neo* [1] | 12 |
| BKK29330 | *trpC2 ΔcmoO::neo* [1] | 12 |
| BKK29340 | *trpC2 ΔtcyN::neo* [1] | 12 |
| BKK29350 | *trpC2 ΔtcyM::neo* [1] | 12 |
| BKK29360 | *trpC2 ΔtcyL::neo* [1] | 12 |
| BKK29390 | *trpC2 ΔsnaA::neo* | 12 |
| BKK29400 | *trpC2 ΔascR::neo* [1] | 12 |

| | | |
|---|---|---|
| BKK30990 | *trpC2 ΔthiT::neo* | 12 |
| BKK32030 | *trpC2 ΔyuiG::neo* | 12 |
| BKK39460 | *trpC2 ΔyxeQ::neo* [1] | 12 |
| BKK39470 | *trpC2 ΔsndB::neo* [1] | 12 |
| BKK39490 | *trpC2 ΔyxeN::neo* [1] | 12 |
| BKK39510 | *trpC2 ΔsnaB::neo* [1] | 12 |
| BKK39520 | *trpC2 ΔyxeK::neo* [1] | 12 |
| GP1171 | *trpC2 xkdE::(N-yfp* $erm^R$*)* | pGP886 → 168 |
| GP3342 | *trpC2 ΔilvC::lox72* | pDR244 → GP4404 (heat cured) |
| GP3343 | *trpC2 ΔpanG::lox72 ΔpanE::lox72* | pDR244 → GP3348 (heat cured) |
| GP3346 | *trpC2 ΔpanG::lox72 ΔilvC::neo* | BKK28290 → GP3347 |
| GP3347 | *trpC2 ΔpanG::lox72* | pDR244 → GP3383 (heat cured) |
| GP3348 | *trpC2 ΔpanG::lox72 ΔpanE::neo* | BKK15110 → GP3347 |
| GP3380 | *trpC2 ΔcoaA::neo* | BKK23760 → 168 |
| GP3381 | *trpC2 ΔcoaX::neo* | BKK00700 → 168 |
| GP3382 | *trpC2 ΔpanD::neo* | BKK22410 → 168 |
| GP3383 | *trpC2 ΔpanG::neo* | BKK14440 → 168 |

| | | |
|---|---|---|
| GP3384 | *trpC2 ΔpanE::neo* | BKK15110 → 168 |
| GP3386 | *trpC2 ΔcysK::neo* | BKK00730 → 168 |
| GP3397 | *trpC2 ΔilvC::lox72 ΔpanE::neo* | GP3384 → GP3342 |
| GP4085 | *trpC2 ΔcymR::neo* | BKK27520 → 168 |
| GP4124 | *trpC2 ΔcysK::lox72 ΔpanB::neo* | GP4401 → GP4462 |
| GP4361 | *trpC2 ΔpanC::cat* | This study |
| GP4362 | *trpC2 ΔpanC::cat ΔcymR::lox72* | GP4361 → GP4463 |
| GP4364 | *trpC2 ΔpanC::cat ΔsnaA::neo* | GP4361 → GP4460 |
| GP4375 | *trpC2 ΔpanC::cat ΔcymR::lox72 ΔtcyN::neo* | BKK29340 → GP4362 |
| GP4378 | *trpC2 ΔpanC::cat ΔtcyN::neo* | BKK29340 → GP4361 |
| GP4379 | *trpC2 ΔpanC::cat xkdE::Pxyl-(empty)-ermC* | GP1171 → GP4361 |
| GP4380 | *trpC2 ΔpanC::cat xkdE::Pxyl-tcyJKLMN-ermC* | pGP4018 (ScaI) → GP4361 |
| GP4383 | *trpC2 ΔpanC::cat ΔsnaA::lox72* | pDR244 → GP4364 (heat cured) |
| GP4401 | *trpC2 ΔpanB::neo* | BKK22430 → 168 |
| GP4402 | *trpC2 ΔpanC::neo* | BKK22420 → 168 |
| GP4403 | *trpC2 ΔpanG::lox72 ΔpanE::lox72 ΔilvC::neo* | BKK28290 → GP3343 |
| GP4404 | *trpC2 ΔilvC::neo* | BKK28290 → 168 |

| | | |
|---|---|---|
| GP4444 | *trpC2 ΔpanC::cat xkdE::Pxyl-tcyJKLMN-ermC ΔcoaA::neo* | GP3380 → GP4380 |
| GP4448 | *trpC2 ΔpanB::neo cysK*$^{G223*}$ *alaS*$^{P731L}$ | GP4401 suppressor on SP (pantothenate depletion) |
| GP4449 | *trpC2 ΔpanB::neo cysK*$^{G223*}$ | GP4401 suppressor on SP (pantothenate depletion) |
| GP4450 | *trpC2 ΔpanC::neo guaB*$^{D64IAMA}$ *[ΔydcL-yddM] yncM*$^{K13E}$ *cymR*$^{L49T}$ | GP4402 suppressor on SP (pantothenate depletion) |
| GP4460 | *trpC2 ΔsnaA::neo* | BKK29390 → 168 |
| GP4462 | *trpC2 ΔcysK::lox72* | pDR244 → GP3386 (heat cured) |
| GP4463 | *trpC2 ΔcymR::lox72* | pDR244 → GP4085 (heat cured) |
| GP4489 | *trpC2 ΔcoaX::neo xkdE::Pxyl-tcyJKLMN-ermC ΔpanC::cat* | GP3381 → GP4380 |
| GP4652 | *trpC2 ΔpanC::cat xkdE::Pxyl-(empty)-ermC ΔcoaA::neo* | GP3380 → GP4379 |
| GP4653 | *trpC2 ΔpanC::cat xkdE::Pxyl-(empty)-ermC ΔcoaX::neo* | GP3381 → GP4379 |
| GP4659 | *trpC2 ΔtcyJKLMN::tet* | This study |
| GP4660 | *trpC2 ΔpanC::cat ΔtcyJKLMN::tet* | GP4659 → GP4361 |

| | | |
|---|---|---|
| GP4670 | *trpC2 ΔcoaBC::spc* | This study |
| GP4694 | *trpC2 ΔpanC::cat ΔribU::neo* | BKK23050 → GP4361 |
| GP4695 | *trpC2 ΔpanC::cat ΔthiT::neo* | BKK30990 → GP4361 |
| GP4696 | *trpC2 ΔpanC::cat ΔtrpP::neo* | BKK10010 → GP4361 |
| GP4697 | *trpC2 ΔpanC::cat ΔpanU::neo* | BKK10370 → GP4361 |
| GP4698 | *trpC2 ΔpanC::cat ΔypdP::neo* | BKK21980 → GP4361 |
| GP4699 | *trpC2 ΔpanC::cat ΔyuiG::neo* | BKK32030 → GP4361 |
| GP4700 | *trpC2 ΔpanC::cat ΔecfT::neo* | BKK01470 → GP4361 |
| GP4703 | *trpC2 ΔecfT::neo* | BKK01470 → 168 |
| GP4704 | *trpC2 ΔpanU::neo* | BKK10370 → 168 |

[1] These mutants were used in the initial screening to elucidate the suppression effect.
